# Stress response, behavior, and development are shaped by transposable element-induced mutations in Drosophila

**DOI:** 10.1101/380618

**Authors:** Gabriel E. Rech, Maria Bogaerts-Marquez, Maite G. Barrón, Miriam Merenciano, José Luis Villanueva-Cañas, Vivien Horváth, Anna-Sophie Fiston-Lavier, Isabelle Luyten, Sandeep Venkataram, Hadi Quesneville, Dmitri A. Petrov, Josefa González

## Abstract

Mapping genotype to phenotype is challenging because of the difficulties in identifying both the traits under selection and the specific genetic variants underlying these traits. Most of the current knowledge of the genetic basis of adaptive evolution is based on the analysis of single nucleotide polymorphisms (SNPs). Despite increasing evidence for their causal role, the contribution of structural variants to adaptive evolution remains largely unexplored. In this work, we analyzed the population frequencies of 1,615 Transposable Element (TE) insertions in 91 samples from 60 worldwide natural populations of *Drosophila melanogaster*. We identified a set of 300 TEs that are present at high population frequencies, and located in genomic regions with high recombination rate, where the efficiency of natural selection is high. The age and the length of these 300 TEs are consistent with relatively young and long insertions reaching high frequencies due to the action of positive selection. Indeed, we, and others, found evidence of selective sweeps and/or population differentiation for 65 of them. The analysis of the genes located nearby these 65 candidate adaptive insertions suggested that the functional response to selection is related with the GO categories of response to stimulus, behavior, and development. We further showed that a subset of the candidate adaptive TEs affect expression of nearby genes, and five of them have already been linked to an ecologically relevant phenotypic effect. Our results provide a more complete understanding of the genetic variation and the fitness-related traits relevant for adaptive evolution. Similar studies should help uncover the importance of TE-induced adaptive mutations in other species as well.

## Introduction

Understanding how organisms adapt to local environmental conditions requires identifying the loci and the phenotypic traits potentially targeted by natural selection, which should also provide critical knowledge for how organisms will respond to environmental change [1-3]. Organisms from plants to humans harbor genetic variation within and among populations that allows them to adapt to diverse local environments [4-6]. Genome scans for selection have almost exclusively focused on identifying single nucleotide polymorphisms (SNPs). However, while the role of other types of genetic variants, such as transposable element (TE) insertions and segmental duplications, in local adaptation has been suggested, these variants are often poorly characterized [7-10]. This is mainly due to technical limitations: short-read sequencing technologies make TE discovery and accurate genotyping difficult. However, deciphering the genetic basis of adaptation requires comprehensive knowledge of these other types of genetic variants, as there is evidence that they are important contributors to adaptive variation [9, 11, 12].

TEs are mobile DNA fragments that constitute a substantial albeit variable proportion of virtually all the genomes analyzed to date [13, 14]. TEs can create a variety of mutations from gene disruption to changes in gene expression and chromosome rearrangements [14, 15]. Although the majority of TE-induced mutations are deleterious or neutral, there are multiple instances in which individual TE insertions have been shown to play a role in adaptive evolution [10-12, 16]. In humans, *MER41* insertions, a family of endogenous retroviruses, have dispersed interferon-inducible enhancers that promote the transcription of innate immunity factors [17]. In *Drosophila melanogaster*, the insertion of an *Accord* retrotransposon in the upstream region of *Cyp6g1* gene leads to transcript up-regulation and increased resistance to several insecticides [18, 19].

However, only a few genome-wide screens have tried to systematically assess the role of TEs in adaptive evolution. In humans, the only screen so far focused on the analysis of a particular TE family, LINE-1 elements, and found that a fraction of these elements showed signatures of positive selection [20]. In *D. melanogaster*, genome-wide screens were initially performed based on a PCR-approach that only allowed studying a subset of all the euchromatic TEs present in the reference genome [7, 8, 21]. In *Arabidopsis thaliana*, genome-wide analysis of TE insertions revealed that TEs affect nearby gene expression and local patterns of DNA methylation, with some of these insertions likely to be involved in adaptation [22, 23]. Thus, while at the moment limited to species with good TE sequence annotations and genome datasets, genome-wide screens for putatively adaptive insertions are a promising strategy to identify genetic variants underlying adaptive evolution [24].

*D. melanogaster* is to date one of the best model systems to identify the genetic and functional basis of adaptive evolution. Originally from sub-tropical Africa, *D. melanogaster* has adapted in recent evolutionary time to a wide-range of environmental conditions [25, 26]. Indeed, there are hundreds of genome sequences available from worldwide populations [27]. This species has one of the best functionally annotated genomes, which facilitates the identification of traits under selection [28]. In addition, TE annotations in the *D. melanogaster* reference genome continue to be updated by the research community [29-31].

In this work, we screened 303 individual genomes, and 83 pooled samples (containing from 30 to 440 chromosomes each) from 60 worldwide natural *D. melanogaster* populations to identify the TE insertions most likely involved in adaptive evolution (Fig 1). In addition to the age and the size of the 1,615 TEs analyzed, we calculated four different statistics to detect potentially adaptive TEs. The GO enrichment analysis of the genes located nearby our set of candidate adaptive insertions pinpoint response to stimulus, behavior, and development as the traits more likely to be shaped by TE-induced mutations. Consistent with these results, genes located nearby our set of candidate adaptive TEs are significantly enriched for previously identified loci underlying stress- and behavior-related traits.

**Fig 1.**
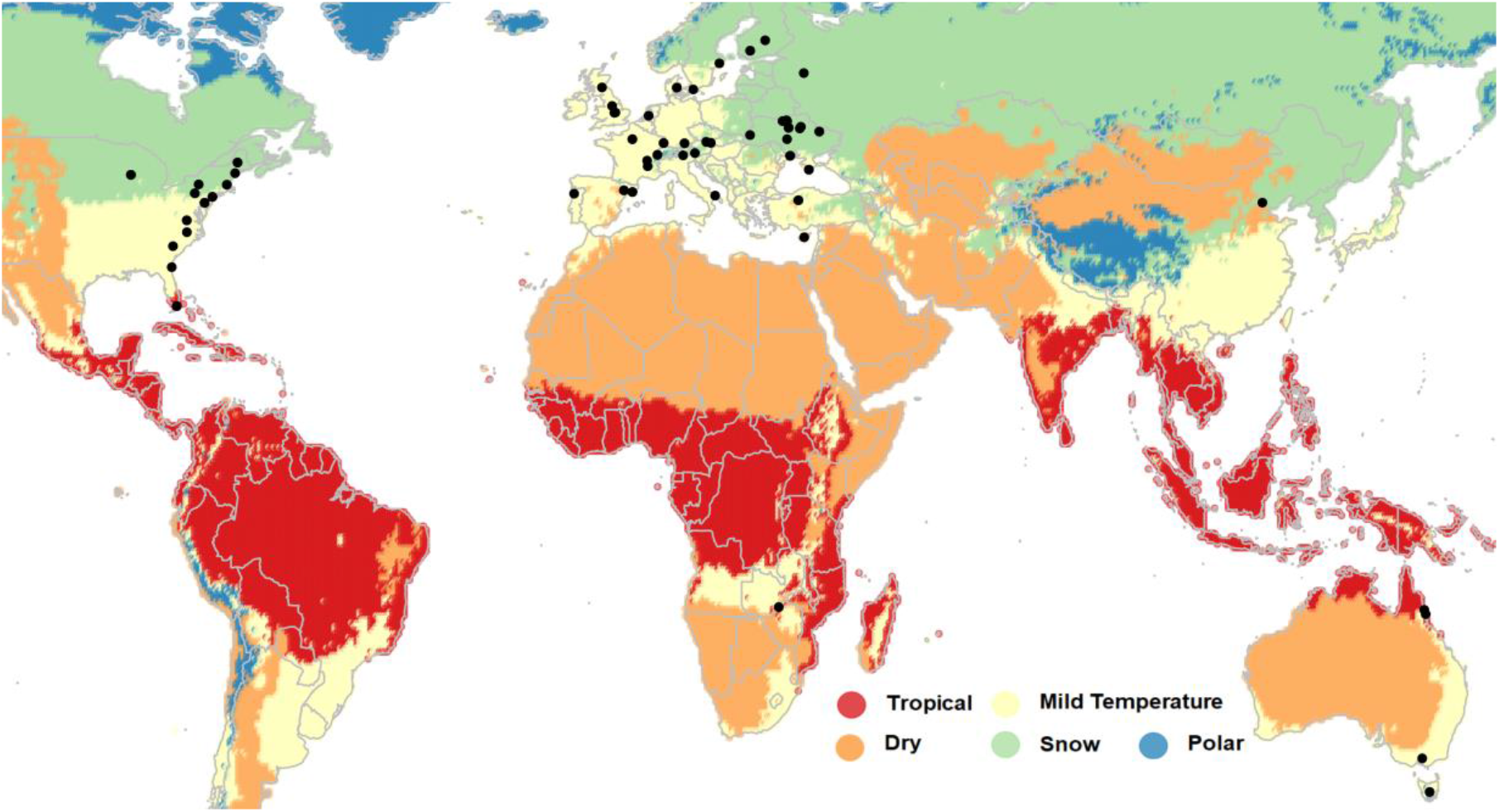
Worldwide distribution of D. melanogaster populations used in this study. Location of the 39 European, 14 North American, five Australian, one Asian, and one African population analyzed in this work. Note that the location of some populations overlap in the map. For more details, see S1 Table. Colors indicate the five major Köppen climate zones [32].

Overall, our results suggest a widespread contribution of TEs to adaptive evolution in *D. melanogaster* and pinpoint relevant traits for adaptation.

## Results

### Natural populations of D. melanogaster contain hundreds of polymorphic TEs at high population frequencies

To identify TEs likely to be involved in adaptation, we looked for TEs present at high population frequencies, and located in genomic regions with high recombination rates (see Material and Methods). We expect TEs that increase the fitness of their carriers to be present at high frequency in the population(s) where adaptation took place [33-36]. In addition, among all the TEs present at high frequencies, TEs located in regions with high recombination rates are less likely to have increased in frequency neutrally compared with TEs located in low recombination regions. This is so because the efficiency of selection in genomic regions with low recombination rates tends to be lower due to the increase in noise generated by linked selection such as background selection and recurrent selective sweeps [37, 38]. Moreover, TEs located in low recombination regions are more likely to be linked to an adaptive mutation rather than being the causal mutation [33-35].

We first estimated population frequencies for 1,615 TE insertions in 91 samples from 60 worldwide natural populations: 39 European, 14 North American, five Australian, one Asian, and one African population collected in the ancestral range of the species (Fig 1 and S1 Table) (see Material and Methods). We classified the 1,615 TEs based on their population frequencies obtained with *Tlex2* [31], and on their genomic location in high or low recombination regions (Fig 2, S2 Table, see Material and Methods). 808 of the 1,615 TEs were present in regions with low recombination rate. Most of these TEs (79%, 640 out of 808 TEs) were fixed, defined here as being present at > 95% frequency in all samples, in all the populations analyzed. Among the 807 TEs located in regions with high recombination rates, 215 were fixed and 177 were present at low frequencies (LowFreq), defined here as being present at ≤ 10% frequency in each of the analyzed samples (Fig 2). Note that the percentage of fixed TEs in high recombination regions is significantly lower than the percentage in low recombination regions (27% vs 79% respectively, Chi-squared p-value = 2.2 ^e-16^), as expected if the efficiency of selection is lower in low recombination regions, and slightly deleterious TEs reached fixation neutrally [37, 38]. Finally, 300 of the 807 TEs located in high recombination regions were present at high frequencies (HighFreq), defined here as being present at < 95% frequency overall and at >10% frequency in at least three samples (Fig 2, S1 Fig).

**Fig 2.**
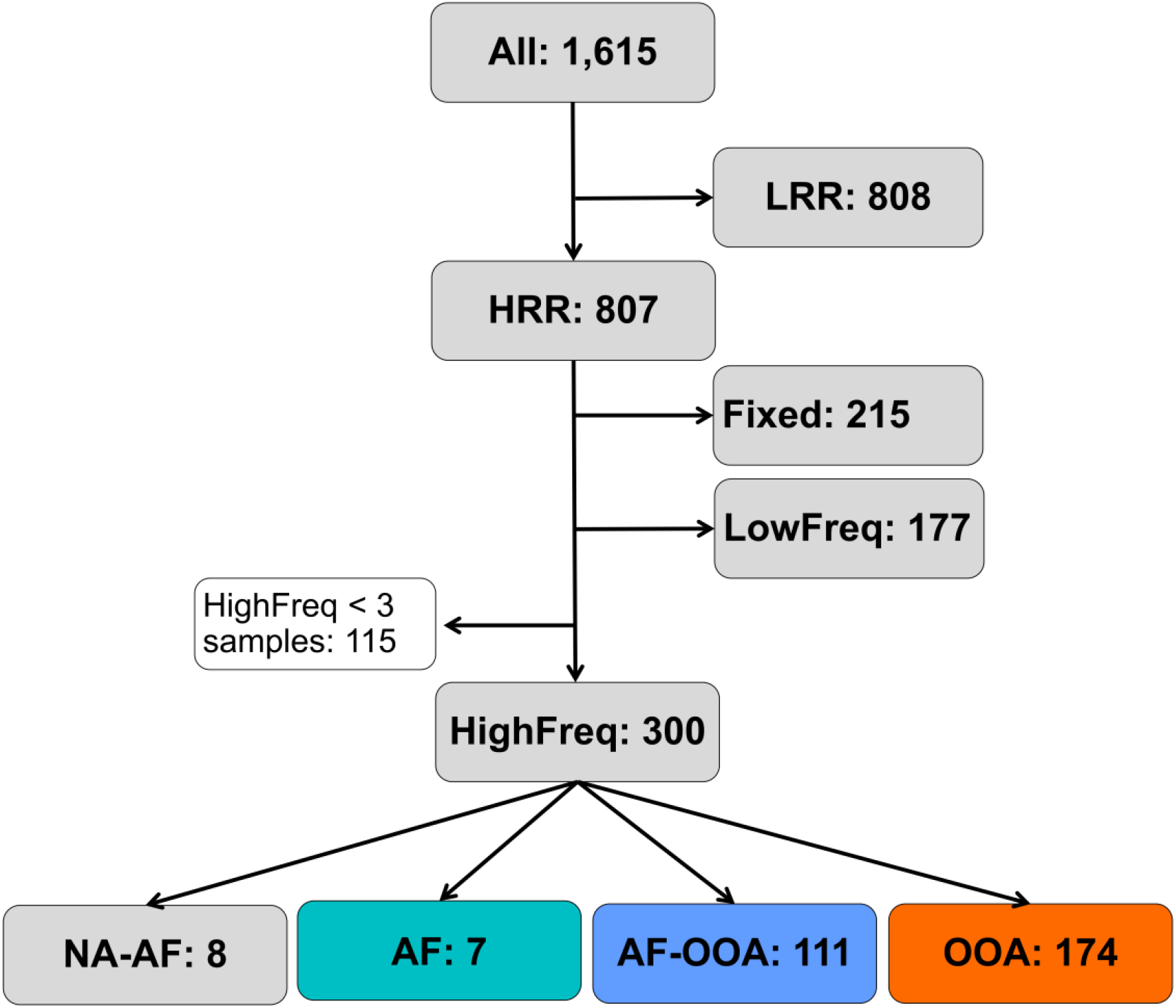
Workflow showing the main steps applied for identifying TEs present at high frequencies in high recombination regions in the D. melanogaster genome. LRR: TEs located at low recombination rate regions. HRR: TEs located at high recombination rate regions. Fixed: HRR TEs at frequencies > 95% in all populations. LowFreq: low frequency HRR TEs (frequencies < 10% in all samples). HighFreq: high frequency HRR TEs (frequencies < 95% in all samples and at >10% frequency in at least three samples). HighFreq TEs were further classified according to their frequency in African (AF) and/or out-of-Africa (OOA) populations: AF: TEs at high frequency only in the African population; AF-OOA: TEs at high frequency in Africa and out-of-Africa populations; OOA: TEs at high frequency in out-of-Africa populations and low frequency in the African population and NA-AF: TEs present at high frequency in out-of-Africa populations but for which we have no data for the African population.

We further classified these 300 TEs according to their frequency in African (AF) and/or out-of-Africa (OOA) populations: seven TEs were only present at high frequencies in the African population analyzed (AF), 111 were present at high frequencies both in African and in the out-of-Africa populations (AF-OOA), and 174 were present at high frequencies only in the out-of-Africa populations (OOA, Fig 2). TEs present at high frequencies both in African and out-of-African populations are more likely to be involved in global adaptations, while TEs present only in African or only in out-of-Africa populations could be involved in local adaptation. Overall, we identified 300 polymorphic TEs present at high frequencies and located in high recombination regions of the genome, which could have increased in frequency due to positive selection. However, it is also possible that some or many of these 300 TEs have increased in frequency neutrally.

### Age and length of TEs present at high frequencies in regions with high recombination are consistent with a putatively adaptive role of these insertions

In addition to the population frequency, the age of a TE insertion can be informative about whether a TE is more likely to be adaptive, neutral, or deleterious. A young TE present at high population frequencies is more likely to have increased in frequency due to recent positive selection, while old TEs present at high population frequencies might have slowly drifted to high frequency [21, 24]. Note that it is entirely possible that such old TEs did increase in frequency due to positive selection and have been maintained by balancing selection since then [39]. Nonetheless, in this paper we primarily focus on the identification of the subset of TEs that are most likely to be adaptive and are willing to tolerate potentially high false negative rates.

We estimated the age of all the TEs annotated in the reference genome using a phylogenetic approach (5,416 TEs, see Material and Methods). We compared our TE age estimates with previously available data for 437 TEs [21, 40]. Among the 417 TEs present in the two datasets, there are 10 TE insertions in our dataset that according to the TE age distributions were outliers (showed much higher age values estimates, S2A Fig). When we removed these 10 data points the correlation between the age estimates from the two studies was high (*r*^2^: 0.71, p-value < 2.2x10^-16^, S2B Fig). Note that the TE age estimates obtained by these methods depend on the dataset used for generating the phylogenies, which differ between the two studies (437 TEs *vs* 5,416 TEs, S2 Fig).

We compared the TE age distributions between the different frequency groups, and we further classified TEs as “young” or “old” insertions according to whether the estimated terminal branch length was < 0.01 or ≥ 0.01, respectively (see Material and Methods). As mentioned above, most of the TEs in low recombination regions are fixed. Accordingly, we found that TEs present in low recombination regions and Fixed TEs in high recombination regions showed similar age distributions (Wilcoxon test, p-value = 0.321, Fig 3A) and contained a large proportion of old TEs, 71% and 75% respectively, as expected if these two datasets contain mostly neutral TEs (Fig 3B, S3 Table). The age distribution of these two groups was different from the LowFreq and the HighFreq groups overall (Wilcoxon test, p-value < 2.2x10^-16^, Fig 3A).

We found that all LowFreq TEs were young TEs (Fig 3B, S3 Table). This result is consistent with LowFreq TEs being slightly deleterious mutations that have not been yet removed from populations by purifying selection. Finally, the three subgroups of HighFreq TEs contained mostly young TEs (Fig 3B, S3 Table).

**Fig 3.**
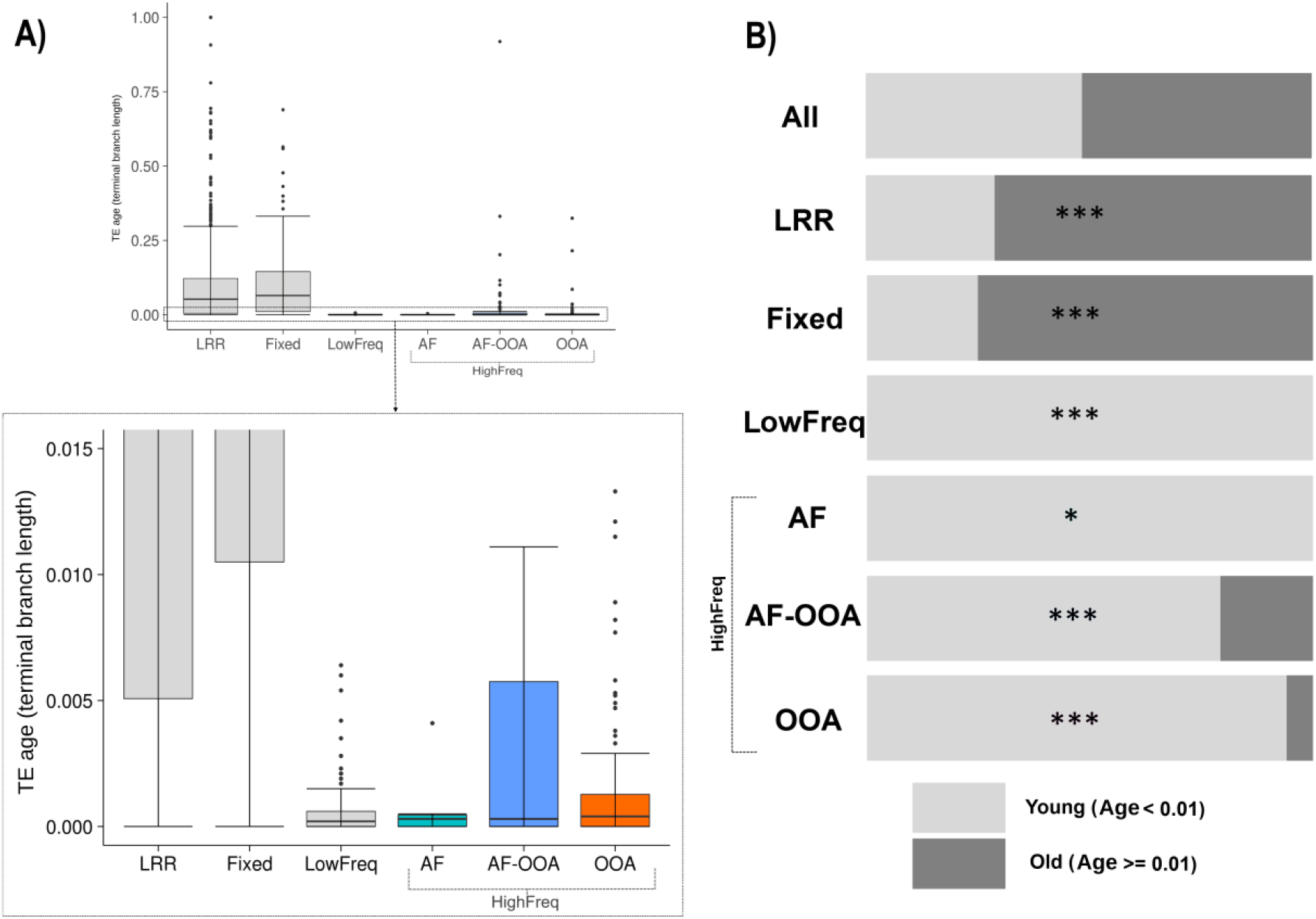
TE age of the different frequency groups. A) Top: Boxplots showing the distribution of TE age (terminal branch length) values for each of the categories. Bottom: Zoomed-in version of the boxed area showing the lowest values of the TE age distribution. B) Proportion of young (age < 0.01) and old (age ≥ 0.01) TEs in each category. * p-value < 0.05, *** p-value < 0.001 from Chi-square test.

The length of a TE can also be informative about whether a TE is more likely to be adaptive, neutral, or deleterious. Because longer TEs are more likely to act as substrates for ectopic recombination leading to deleterious rearrangements, if a TE is long but it is present at high population frequencies, it is more likely to be adaptive [16, 41, 42]. In contrast, shorter TEs are both more likely to be nearly neutral in their selective effect due to lower rate of ectopic recombination among shorter homologous sequences and in addition more likely to be older and thus shorter because of the high rate of DNA loss in Drosophila [43]. We used the TE length ratio, calculated as the proportion of the length of the TE insertion regarding the length of the canonical family sequence, as a proxy for measuring the relative length of the TEs in each group. We found statistically significant differences between the HighFreq and the other three TE groups: LowFreq, Fixed, and TEs in low recombination regions (S4 Table). In particular, HighFreq and LowFreq TEs show distributions of TE Length Ratio shifted upwards (median: 59.3 and 80.4 respectively), while the distributions of Fixed TEs and TEs in low recombination regions are shifted downwards, showing a predominance of shorter TEs (mean: 16.2 and 30.7 respectively) (Fig 4 and S4 Table). No differences in the TE length ratio among the HighFreq TEs subgroups were found (Kruskal Wallis test, p = 0.062) (S4 Table).

**Fig 4.**
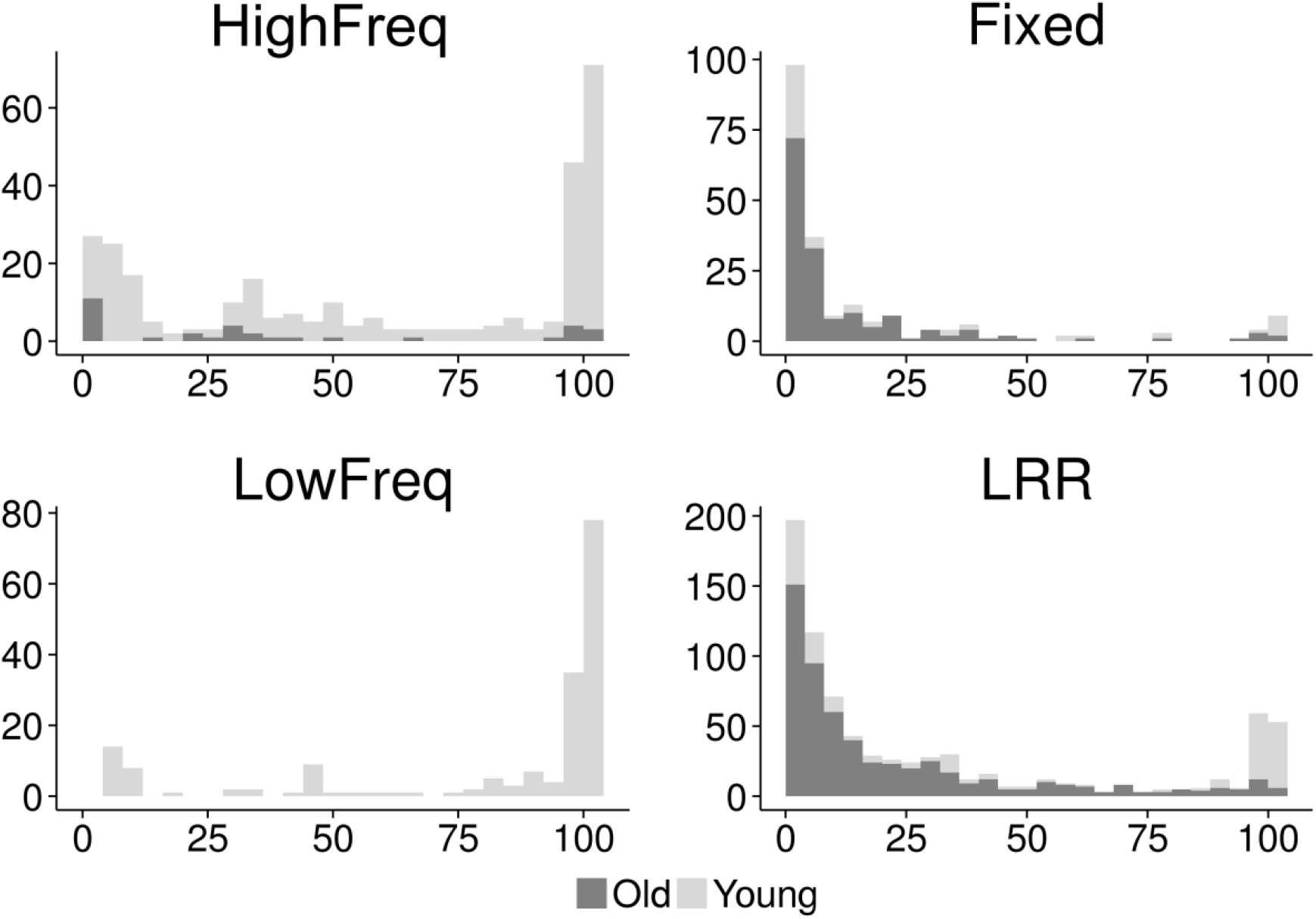
Number of TEs at different TE Length Ratios (%). Bars indicate number of TEs (vertical axis) per bin of TE Length Ratio (%) (horizontal axis) and color shade indicates the proportion of young and old TEs in each bin.

When considering both age and length of the TEs across different categories, we found that Fixed TEs and TEs in low recombination regions show predominance of older and truncated TEs (Fig 4), which is consistent with old TE insertions that have reached fixation through processes other than positive selection. On the other hand, the HighFreq and LowFreq groups contain mostly large and young TEs (Fig 4). In the case of LowFreq TEs, these results are consistent with the hypothesis that low frequency TEs could be recent insertions that purifying selection still did not have time to eliminate. Finally, young and large HighFreq TEs support the hypothesis of the presence in this group of a large number of recent putatively functional insertions that have rapidly increase in frequency due to the action of positive selection. Thus, for the rest of this work, we focused on HighFreq TEs to look for further evidence suggesting their contribution to adaptive evolution.

### TEs present at high frequencies in high recombination regions showed different signatures of positive selection

To test whether HighFreq TEs showed signatures of positive selection, we used two different approaches: we looked for signatures of selective sweeps in the regions flanking the candidate adaptive TEs, and we looked for evidence of population differentiation between populations located at the extremes of latitudinal clines in three continents: Europe (EU), North America (NA), and Australia.

To look for signatures of selective sweeps in the vicinity of the candidate TE insertions, we used three different haplotype-based methods in order to identify different signals of selective sweeps: (i) the *iHS* test mainly detects events of hard sweeps [44], (ii) the *H12* test detects both hard and soft sweeps [45], and (iii) the *nS*_L_ test detects sweeps under different scenarios, and it is more robust to recombination rate variation [46]. We independently applied these tests to two datasets: one dataset containing 141 strains from the Raleigh population in NA, and a second dataset containing 158 strains from four different populations in EU. Note that EU populations do not show latitudinal population structure, and thus we analyzed them together [47] (see Material and Methods). Overall, we were able to calculate at least one test, in at least one of the two continents, for 202 of the 300 HighFreq TE insertions (S5 Table). To determine the significance of iHS and *nS*_L_ values, we compared them with the distribution of values obtained from neutral SNPs, while for H12 we selected the top 15% values (see Material and Methods). Overall, 36 TEs showed evidence of selection (Fig 5 and S6 Table). The three tests identified similar numbers of significant TEs (Chi-square test, p-value = 0.350, S5 Table), however the overlap between the TEs identified by the different tests was low (S3A Fig). These results suggest that these 36 TEs could be evolving under different selective scenarios, including both hard and soft sweeps.

We also tested whether the signals of selection differ among continents. For 31 out of the 36 TEs that showed signatures of selection we had data from NA and EU populations. However, only 6 of these 31 TEs showed evidence of selection in both continents while the other 25 TEs were significant only in NA or only in EU, suggesting that the signatures of selection could be continent specific (S3B Fig). Finally, while *iHS* and *nSL* identified similar numbers of TEs in the two continents, H12 identified more significant TEs in NA (Chi-square test, p-value = 0.032, S5 Table).

**Fig 5.**
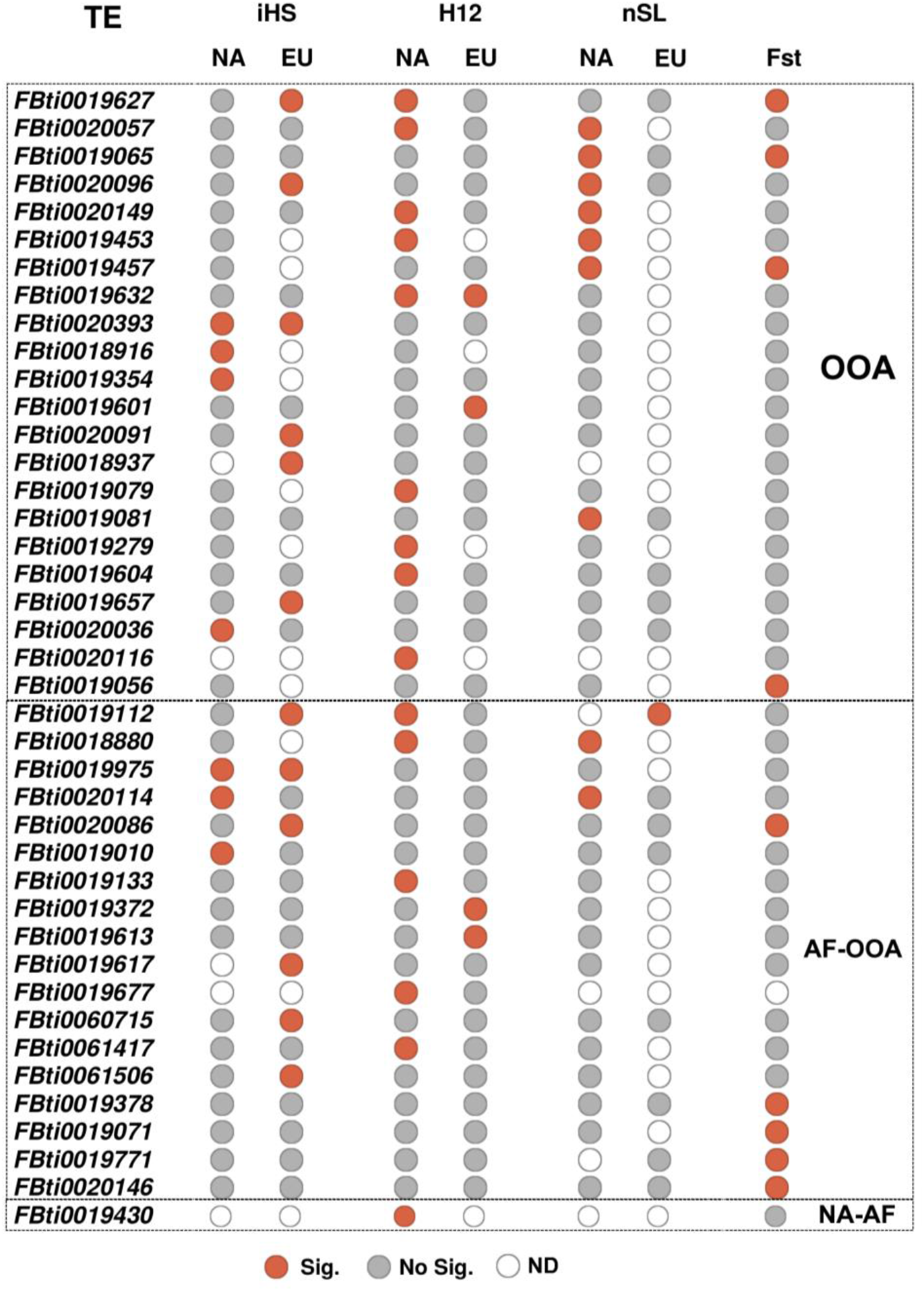
HighFreq TEs with signals of selection. 41 HighFreq TEs showing at least one signal of selection either or both in the selective sweep tests (iHS, H12 or *nS*_L_, 36 TEs) or the population differentiation test (Fst, 9 TEs). Red and grey circles indicate statistical significance for each TE at each test and population (Significant and No significant, respectively). Empty circles (ND) indicates that the test could not be calculated.

Besides selective sweeps, we also looked for evidence of population differentiation using the pairwise FST estimator of Weir & Cockerham (1984) [48]. We performed six pairwise comparisons among latitudinal distant populations: two populations in EU, two in NA, and two in Australia (see Materials and Methods). We could estimate FST for 254 of the 300 HighFreq TE insertions (S7 Table). To determine the significance of FST values, we compared them with the distribution of values obtained from neutral SNPs in each pair of populations (see Material and Methods). 78 TEs show significant FST values, and we further filter them by keeping only those that were significant in more than one pairwise comparison and consistently present at high frequencies in populations located in high latitudes or in low latitudes (concordant FST) (see Material and Methods). After this filtering step, nine TEs were significant (S4 Fig). Five of these nine TEs were also identified as being under positive selection by at least one haplotype-based test (Fig 5). Overall, we could calculate at least one statistic for 273 HighFreq TEs, and 41 of them showed evidence of positive selection (Fig 5, S5 Table). TEs present at high frequencies both in African and in the out-of-Africa populations (AF-OOA), and TEs present at high frequencies only in the out-of-Africa populations (OOA) showed similar percentage of TEs with evidence of selection, 18/103 (17.5%) and 22/154 (14.2%) respectively (Chi-square, p-value = 0.488, S5 Table), suggesting that both datasets could be enriched for adaptive TEs. Indeed, 10 of these 41 TEs were previously found to show evidence of positive selection (Table 1).

**Table 1.**
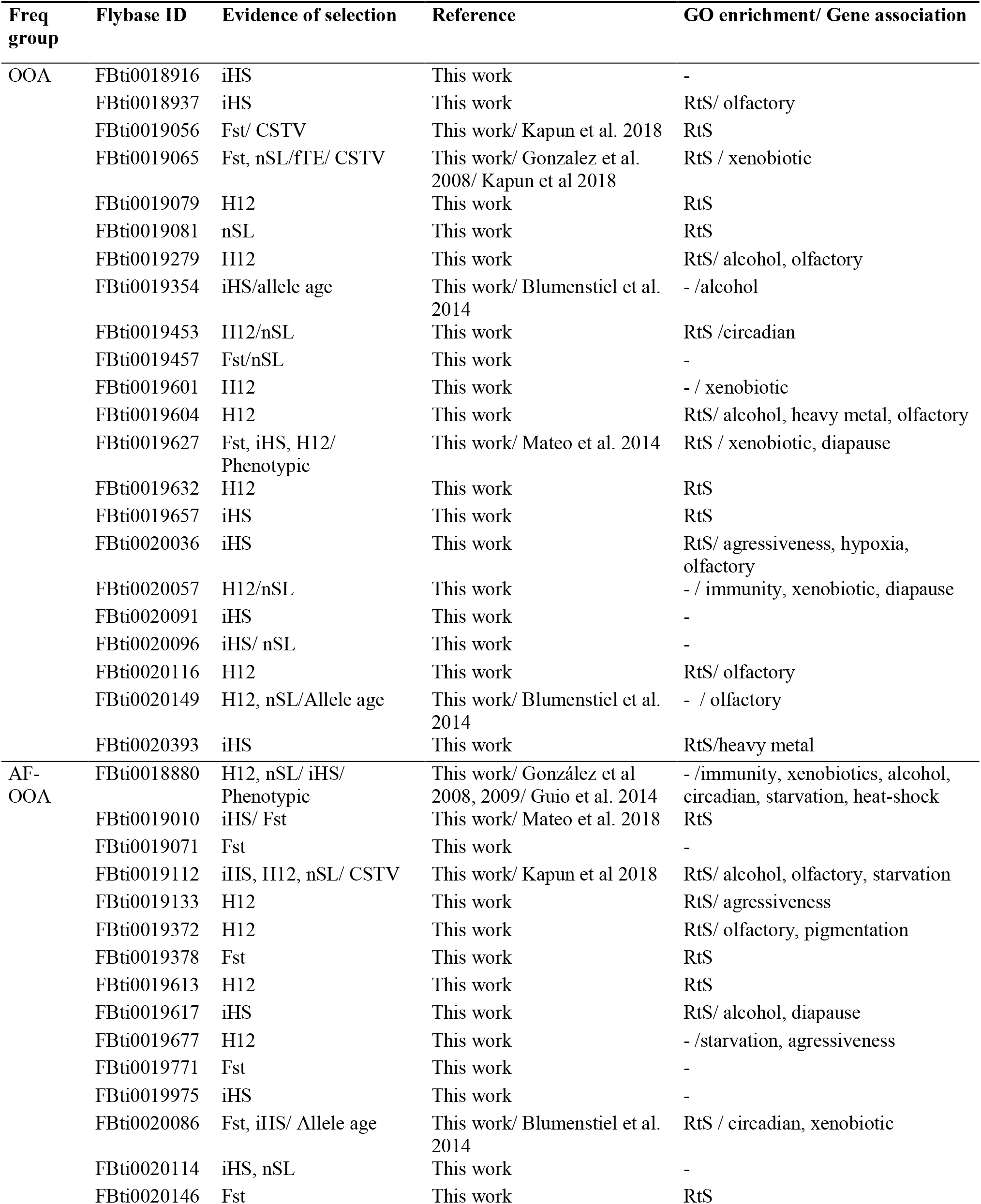
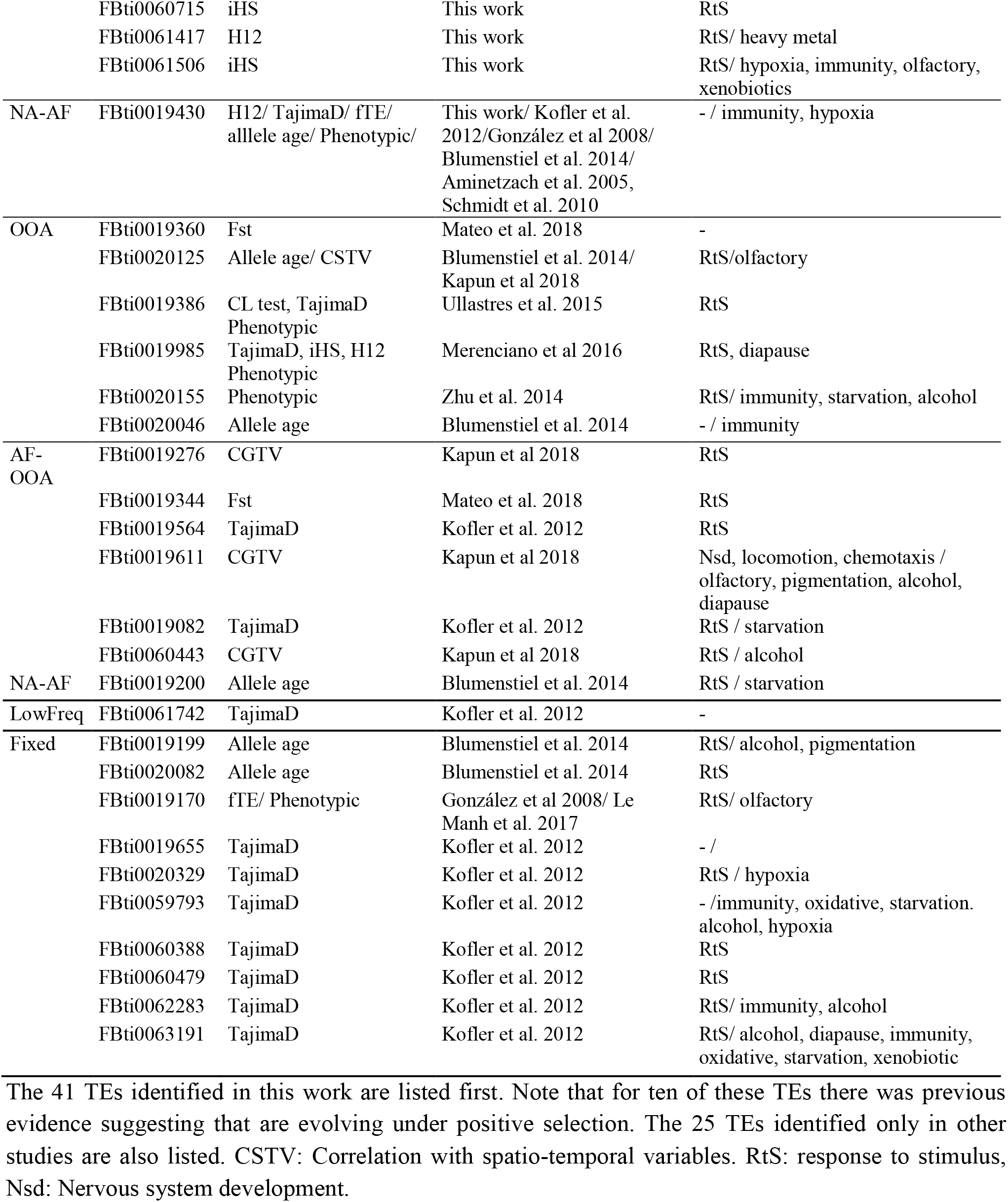
65 TEs showing evidence of selection (ES).

### Candidate adaptive TEs are associated with genes involved in stress response, behavior, and development

We used the GO terms of genes nearby candidate adaptive TEs to test whether they were enriched for any biological processes. Besides, the 41 TEs identified in this work, we also consider 24 TEs that have been previously identified as candidate adaptive TEs based on different approaches such as Tajima’s D, and age of allele neutrality test (Table 1). In total, we analyzed 83 genes nearby 65 TEs (Table 1, S8A Table). We found four significant clusters (enrichment score > 1.3) according to DAVID [49, 50] functional annotation tool: response to stimulus, behavior, development, and localization and transport (Fig 6A, S8A Table). We then analyzed whether the 363 genes nearby the 300 HighFreq TEs were enriched for similar biological processes (see Material and Methods). We identified 20 significant clusters (S8A Table). Among clusters showing the highest enrichment scores we also found GO terms related with response to stimulus, behavior and learning, and development (Fig 6B). Finally, genes nearby OOA and AF-OOA TEs were also enriched for similar biological functions (S5 Fig, S8C-D Tables). Note that the behavior-related clusters slightly differed among the datasets: genes nearby TEs with evidence of positive selection were enriched for aggressiveness genes, genes nearby HighFreq TEs and AF-OOA TEs were enriched for olfactory genes, and genes nearby OOA TEs for circadian and locomotor behavior genes (Fig 6 and S5 Fig).

**Fig 6.**
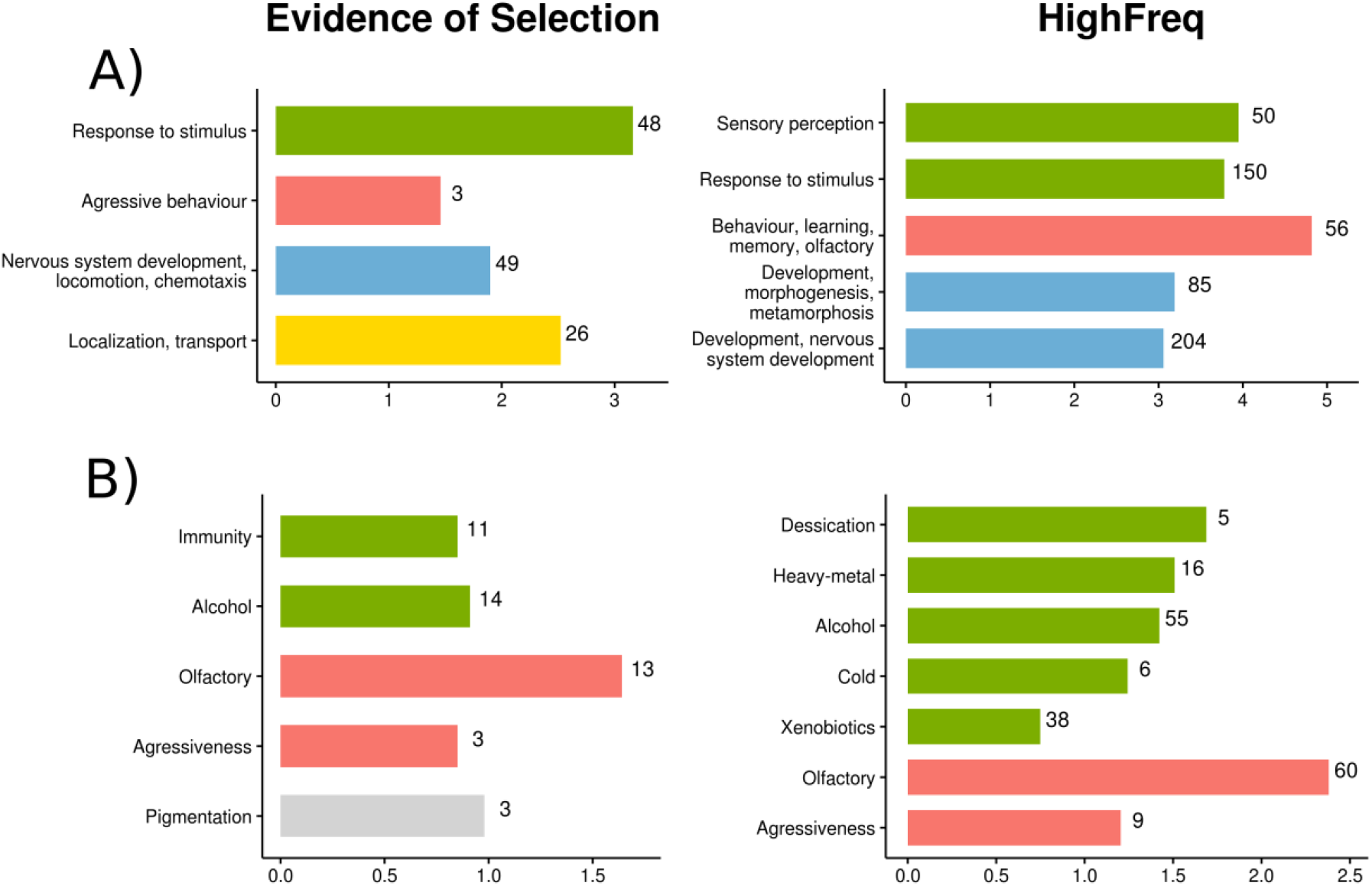
Functional Enrichment analysis of genes nearby TEs showing Evidence of Selection (in this or previous works) and HighFreq TEs. Bar colors indicates similar biological functions of the DAVID clusters (A) and the fitness-related traits (B): Green: stress response, Red: behavior, Blue: development Yellow: transport, Grey: pigmentation. A) Significant gene ontology clusters according to DAVID functional annotation tool (enrichment score > 1.3). For genes nearby HighFreq TEs, only top five clusters are showed. The horizontal axis represent DAVID enrichment score (see S8A and S8B Tables for details). B) Significantly overrepresented fitness-related genes according to previous genome association studies. All FDR corrected p-values < 0.05, Chi-square (χ^2^) test (see S10A and S10B Tables for details). The horizontal axis represents the log10(χ^2^). In both, A) and B), numbers nearby each bar indicate total number of genes in that cluster/category.

To gain more insight into the function of genes nearby the candidate adaptive TEs, we looked whether they were previously described as candidate genes for several fitness-related traits (S9 Table, see Material and Methods). Among the 83 genes nearby the 65 candidate adaptive TEs, 19 have previously been identified as candidates for stress-related phenotypes: 11 genes were associated with immunity and 14 with alcohol exposure (Fig 6B, S10A Table). In addition, we also found enrichment of genes related with behavioral phenotypes such as olfaction and aggressiveness, and with pigmentation (Fig 6B, S10A Table). Similar enrichments were found for genes located nearby the 300 High Freq TEs and for the genes located nearby the OOA and the AF-OOA datasets (S5 Fig, S10C-D Tables). Among the 363 genes nearby HighFreq TEs, 171 have previously been identified as candidates for stress-related phenotypes, such as desiccation, heavy-metal and alcohol, and/or behavior-related phenotypes (Fig 6B).

Overall, we found that genes nearby the 300 HighFreq TEs are enriched for similar biological processes as genes nearby a dataset of TEs with evidence of positive selection: response to stimulus, behavior and learning, and development Fig 6A, S8 Table). Moreover, 47% of the genes nearby the 300 HighFreq TE dataset have previously been identified as candidate genes for several stress- and/or behavior-related traits (Fig 6B, S10 Table).

### Candidate adaptive TEs correlate with the expression of nearby genes

We tested whether there was a correlation between the presence of the candidate adaptive TEs and the expression of nearby genes using the *Matrix eQTL* package [51]. We used gene expression data from Huang *et al*. [52] and *T-lex2* annotations for 140 DGRP lines in order to determine whether the presence of a TE was correlated with the expression level of the nearby genes (< 1kb). We calculated correlations for 638 TEs located at high recombination regions and we found that 19 of them showed significant eQTL associations (S11 Table). TEs present at high frequencies contained more significant eQTLs than expected (38% vs 11%, Chi-Square test, p-value < 0.0001) (Table 2). We observed the same significant tendency when considering only positive correlations (the presence of the TE correlates with increased expression of the nearby gene) or only negative correlations (the presence of the TE correlates with reduced expression of the nearby gene) (Table 2). These results remained significant after FDR correction (50% vs 11% expected, Chi-Square test, p-value < 0.0001, Table 2). Of the 19 TEs showing significant eQTL associations, 11 also showed signatures of selection (S11 Table).

**Table 2.** Correlation between TEs and expression level of nearby genes.

| TEs |  | HighFreq |  | Fixed |  | LowFreq |  | Private |  |
| --- | --- | --- | --- | --- | --- | --- | --- | --- | --- |
| **All TEs analyzed** |  | 70 | 11% | 192 | 30% | 376 | 59% | 25 | 4% |
| **Significant TEs** | **All** | 19 | 38% (**) | 12 | 24% | 19 | 38% | 4 | 8% |
|  | **Positive correlation** | 15 | 37% (**) | 11 | 27% | 15 | 37% | 4 | 10% (*) |
|  | **Negative correlation** | 11 | 32% (**) | 8 | 24% | 15 | 44% | 3 | 9% (*) |
|  | **FDR<0.05** | 5 | 50% (**) | 0 | 0% | 5 | 50% | 0 | 0% |

Number of TEs located at high recombination regions for which correlations were calculated (All TEs analyzed), and number of TEs with significant correlations for each frequency group are given (Significant TEs). Frequency groups were determined based on their frequency in the DGRP population. LowFreq TEs were further classified as Private if only one strain was containing the TE. Note that TEs are classified as fixed if they are present in > 95% of the strains analyzed, thus for some of these TEs there could be strains that do not contain the insertion. Percentages regarding the total number of TEs in that frequency category are also given. Chi-square test \* p-value < 0.05 and \*\* p-value < 0.0001.

We finally checked whether private TEs (those present in only one DGRP strain according to *T-lex2*) were also present among the significant eQTL as expected by the “rare alleles of large effect” hypothesis [53]. We found a small, but still significant set of private TEs with significant correlation with the expression of nearby genes (10% and 9% vs 4% expected, Chi-Square test, p-value < 0.050) (Table 2), which is in agreement with previous reports [54].

### Genomic location, order, and family enrichment of TEs present at high frequencies in high recombination regions

We tested whether the genomic location of HighFreq TEs differed from the location of all TEs in the genome. We classified the TEs as present in intergenic, promoter, or genic regions (see Material and Methods). We found no differences between the distributions of HighFreq vs all TEs in the genome (Chi-square test, p-values > 0.05, Fig 7A, S12A Table). Similar results were obtained when we considered the three HighFreq TEs subgroups (S12A Table). We further classified intragenic TEs in exonic, UTRs, 1^st^ intron, and other introns. Only HighFreq TEs were enriched in UTR regions (Chi-square test, p-value < 0.043) (Fig 7B, 12B Table).

**Fig 7.**
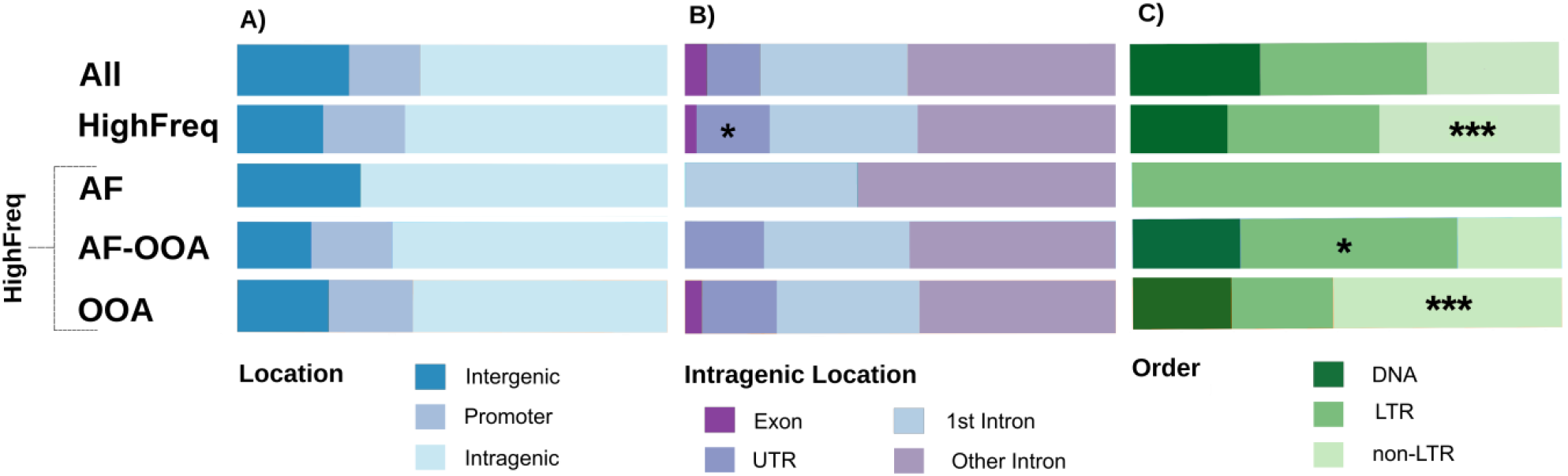
Caracteristics of the HighFreq TEs. **A**) TE location regarding the nearest gene. **B**) Location of intragenic TEs. **C**) TE order. *: p-value < 0.05. ***: p-value < 0.001 (Chi-square test).

We also checked whether the proportion of DNA, LTR, and nonLTR TE orders differed between HighFreq TEs and all TEs in the genome. We found that the HighFreq group contains a larger proportion of non-LTR TEs (42% vs 31% and 33%, Chi-square test, p-value = 5.73e-06, Fig 7C, S13 Table). Moreover, when considering HighFreq subgroups we found that OOA TEs also contain a large proportion of non-LTR elements (53% vs 31% and 33%, Chi-square test, p-value = 1.79e-11) while the AF-OOA TEs contain more LTR elements (50% vs 39%, Chi-square test, p-value = 1.08e-02) (Fig 7C, S13 Table).

Regarding TE families, we found that the HighFreq TEs contain a larger proportion of several families including *jockey, 297, BS* and *pogo* families (S14 Table). When considering only OOA TEs, we found a larger proportion of several families including *jockey, F* family, and *BS*, while in the AF-OOA there was a larger proportion of *297, Quasimodo*, and *opus* (Chi-square test, Bonferroni corrected p-values < 0.05) (S14 Table).

## Discussion

In this work, we identified 300 TEs present at high frequencies in natural populations, and located in genomic regions with high recombination, where the efficiency of selection is high [37, 38]. Most of these TEs are young insertions suggesting that they have increased in frequency relatively fast (Fig 3). In addition, these insertions are longer compared with other TEs in the genome, also suggesting an adaptive role because long insertions are more likely to act as substrates for ectopic recombination leading to chromosome rearrangements that are often deleterious [16, 41, 42] (Fig 4). Our dataset of 300 putatively adaptive TEs, contains all the insertions present at high population frequencies that have previously been identified as putatively adaptive [7, 21, 55-63]. Note that we, and others, have found signatures of positive selection and/or functional evidence for the adaptive role of 53 of the 300 putatively adaptive TEs identified in this work, further suggesting that this dataset is enriched for adaptive insertions (Table 1). The other 12 TEs that have been previously identify as candidate adaptive TEs were fixed or present at low frequencies in the populations analyzed in this study, and thus were not included in our dataset of high frequent TEs (Table 1).

Although we looked for evidence of hard and soft sweeps, and for evidence of population differentiation using the FST statistic, adaptive mutations could show other signatures of selection as well [1, 2, 64]. Polygenic adaptation, which should lead to modest changes in allele frequency at many loci, would be overlooked by the more conventional methods for detecting selection used in this study [65]. A recent work used FST and gene set enrichment analysis to find evidence of polygenic adaptation in European *D. melanogaster* populations [63]. In addition, analysis of environmental correlations between allele frequencies and ecological variables could also lead to the identification of additional TE insertions under positive selection [66-69]. Thus, further analysis could lead to the identification of signatures of selection in other insertions in our dataset besides the 53 insertions that showed signatures of selection identified in this work (Table 1).

Our dataset of 300 putatively adaptive TEs allowed us investigating global patterns in the biological functions that might be affected by TE-induced adaptive mutations in the *D. melanogaster* genome. Previous genome-wide screenings looking for adaptive TE insertions identified a small number of candidates that preclude the identification of the putative traits under selection [7, 8, 21, 61]. In this work, we found that genes nearby putatively adaptive TEs are enriched for response to stimulus, development, and behavioral and learning functions (Fig 6). Through literature searches, we found that 41% (148 out of 363) of these genes have previously been identified as candidate stress-related genes including xenobiotic stress, desiccation, and cold stress (Fig 6). If we focus on the subset of TEs that are likely to be involved in out-of-Africa adaptations, we found similar gene functional enrichments (S5 Fig). Interestingly, circadian behavior gene functions are enriched in this dataset of TEs, consistent with adaptation to seasonal changes in daylight experienced by flies in their out-of-Africa expansion [70]. Thus, our results showed that TE-induce adaptive mutations are mainly likely to contribute to stress-response, developmental, and behavioral traits. Although these traits have previously been identified as targets of natural selection, our results point to the most likely causal variant rather than to a group of linked SNPs [71-73]. Thus, although challenging and time-consuming, follow-up functional analysis of these adaptive mutations should confirm their causal role, as we, and others, have already demonstrated in the past [55-60, 62].

Most of the signatures of positive selection found in the regions flanking the putatively adaptive insertions were continent specific (Fig S3B). These results suggest that a significant proportion of the 300 putatively adaptive TEs could be involved in local adaptation. Thus, it is likely that by exploring more natural populations we could identify additional adaptive insertions. We are also missing TEs that could be playing a role in seasonal and altitudinal adaptation, as both dimensions have been shown to be relevant for *D. melanogaster* [74-76]. Finally, our study is also limited to those insertions present in the reference genome. Although there are several packages that infer the presence of *de novo* TE insertions in genome sequencing data, none of them provides the precise genomic coordinates of the insertions, which result in inaccurate TE frequency estimations [10, 77]. In addition, the size and the age of the *de novo* insertions cannot be estimated hindering the characterization of putatively adaptive insertions [77, 78]. Long-read sequencing techniques should, in the near future, help overcome this limitation and allow the community to investigate the contribution of non-reference TE insertions to adaptive evolution [79].

We also found that the presence of 19 of the candidate adaptive TEs correlated with changes in expression, both up-regulation and down-regulation, of nearby genes (Table 2 and S11 Table). For four of these TEs, *FBti0018880, FBti0019627, FBti0019386*, and *FBti0019985*, changes in expression of the nearby genes have also been reported based on allele-specific expression and/or qRT-PCR experiments, and further shown to be associated with changes in fitness-related traits [56-59, 80]. In addition to these 19 insertions, another four TEs *FBti0020119, FBti0020057, FBti0018883*, and *FBti0020137* were associated with allele-specific expression changes [80]. Thus, overall, 23 insertions are associated with changes of expression of nearby genes, which at least in four cases lead to changes in fitness-related traits. Note that because 41% of the genes nearby candidate adaptive TEs are candidates for stress-related phenotypes, it could be that changes in expression are only induced by the TEs in response to stress.

Overall, we identified 300 TE insertions likely to be involved in adaptive evolution as suggested by their population frequencies, age, size, and presence of signatures of selection in a subset of them. These TEs substantially add to the list of genetic variants likely to play a role in adaptation in *D. melanogaster*. Functional profiling of these candidates should help elucidate the molecular mechanisms underlying these mutations, and confirm their adaptive effect on the traits identified.

## Material and Methods

### Dataset

We analyzed available *D. melanogaster* genome sequencing datasets from 91 samples collected in 60 natural populations distributed worldwide (Fig 1 and S1 Table). Most samples (83) were generated using pool-sequencing, while the remaining eight samples came from individually sequenced strains. The distribution of populations across continents was: one from Asia, 39 from Europe, 14 from North America, five from Oceania, and one from Africa. The African population was collected in Zambia, the ancestral range of the species [81]. For this work, we only used the 67 Zambian strains without any European admixture [81]. All data was downloaded from the NCBI Sequence Read Archive (SRA) from published projects as of April 2016, and from data available in our laboratory (S1 Table). Note that we attempted to include five more samples in our dataset, but we were unable to estimate TE frequencies in these samples. These samples were from Queensland and Tasmania [71], Winters [82], Vienna [83], and Povoa de Varzim [61].

### Transposable element frequency estimation

To estimate TE population frequencies, we used *T-lex2*, a computational tool that works both with individual genomes and with pooled samples. *T-lex2* combines the genotyping information obtained for each individual genome to calculate the population frequency, while for pooled samples the frequency is directly estimated from the number of reads providing evidence for the presence and for the absence of each insertion [31]. Population frequencies for 34 European populations estimated using *Tlex-2* were obtained from Mateo *et al*.[63] and Kapun *et al*.[47]. We used *T-lex2* [31] to estimate the population frequency in the other 26 available populations (six populations sequenced as individual genomes, and 20 populations sequenced as pooled samples).We first downloaded genomic coordinates of all the annotated TEs (5,416 TEs) from FlyBase r6.04 [84, 85]. 2,234 of the 5,416 TEs belong to the INE family that has been inactive for the past 3-4.6 Myr [86], and were discarded. From the 3,182 non-INE TEs, we excluded nested TEs, TEs flanked by other non-INE TEs (100bp on each side of the TE), and TEs that are part of segmental duplications, because *T-lex2* does not provide accurate frequency estimates for these TEs [31]. After these filtering steps we end up with 1,630 TEs. For 108 of the 1,630 TEs we used the corrected genomic coordinates as described by Fiston-Lavier *et al*. [31]. *T-lex2* parameters were set to default except for read length and the use of paired reads that were specific to each dataset.

For the eight individually-sequenced populations, *T-lex2* was able to calculate frequencies for the 1,630 TEs in most of the strains (S6 Fig). Indeed, we only considered a TE frequency if we had data from at least 9 strains in a given population, as this is the smallest number of strains in a sample (S6 Fig). For the 83 samples that were pool-sequenced, we only considered frequencies calculated with 3 to 90 reads. These minimum and maximum thresholds were selected after comparing the distribution of reads in the 48 DrosEU samples to avoid false positives (very low number of reads) or an excess of coverage due to non-unique mapping or spurious reads [47] (S7 Fig). For one population, we have both individually sequenced genomes, and pooled-sequenced genomes. Using data of the individually sequenced population of Stockholm [63] we found a high correlation with the pool-sequenced data of the same population (Pearson correlation coefficient r=0.98, p-value < 2.2e-16, S8 Fig), which indicates that there is no bias due to the sequencing strategy when calculating the frequencies using *T-lex2*. For most TEs we could estimate frequency in most of the samples (S9 Fig). We only discarded 15 TEs where *T-lex2* estimated frequencies for less than 10 out of the 91 samples, ending up with a dataset of 1,615 TEs.

We considered a TE to be located in high recombination regions when the two available recombination estimations for *D. melanogaster* [87, 88] were greater than 0 in the region were the TE is inserted (S2 Table).

### Detecting inversions and correcting TE frequencies

We analyzed the effect of inversions in TE frequency estimations. We focused on the cosmopolitan inversions: In(2L)t, In(2R)Ns, In(3L)P, In(3R)K, In(3R)Mo, In(3R)Payne, and In(3R)C (S15 Table) [75]. 358 TEs are located inside or overlapping with one of these inversions and 36 TEs are located less than 500kb from an inversion breakpoint. For five samples, there is data available on the presence/absence data of inversions: Zambia [81], France [89], North Carolina (DGRP, USA) [90, 91], Italy and Sweden [63]. For all these datasets, we re-estimated TE frequencies for individual samples by removing the strains containing an inversion. We also removed strains where a TE was located 500 kb upstream or downstream of an inversion present in that strain [75]. Removal of strains was done at the TE level using an *in house* python script. As a result, each TE had a different number of supporting strains. The frequencies calculated removing strains with inversions were equivalent to the original ones (Pearson correlation coefficient r = 0.99, p < 2.2e-16, S10 Fig), indicating that the effect of inversions on TE frequency is rather small in our dataset.

### TE age and TE length ratio

We used a phylogeny-based approach to estimate the age of each TE within each family for the 5,416 TEs annotated in the reference genome. The age was estimated as the unique number of substitutions shared between the two closest TEs assuming that they all derived from a common ancestral TE, *i.e*. the divergence between closest TEs. Hence, this approach estimates the time since last activity for each TE. Note that activity includes not only transposition but also other genomic TE movements such as the ones caused by duplications.

When the age estimates were calculated, TE annotations were only available for the release 4. Thus, we started by detecting and annotating the TE families and subfamilies in the release 5 of the reference *D. melanogaster* genome. We used the *de novo* homology based approach developed in the REPET suite to build a library of TE consensus [92] (https://urgi.versailles.inra.fr/Tools/REPET/). The consensus are proxies of the TE family and subfamily canonical sequences. We then annotated each consensus by blasting them against the TE canonical sequences from the Berkeley Drosophila Genome Project (www.fruitfly.org/). Each TE sequence was then aligned to its set of annotated TE consensus using a global alignment tool from the REPET suite, called RefAlign. The RefAlign launches pairwise alignments avoiding spurious alignments induced by internally deleted TE sequences [30, 93]. All pairwise alignments from the same TE family were re-aligned to generate profiles using Clustalw v2.0.10 [94]. We manually curated each profile: we removed shared substitutions and indels using another tool in the REPET suite called *cleanMultipleAlign.py* [30, 93]. A limitation of alignment-based methods is that short TEs could generate misalignments. Hence, to reduce the impact of misalignments 25 TEs shorter than 100bp were removed. For eight TE families (*aurora, BS4, frogger, R1-2, Stalker3*, *TART-B*, *TART-C*, and *Xanthias*) composed by less than three copies, we failed to estimate the divergence of the copies and were not considered in this study (11 copies in total). Some profiles were re-aligned using MAFFT v.7 in order to refine conserved regions between TE sequences [95]. For each TE profile, a phylogenetic tree was inferred using the phyML program with the Hasegawa–Kishono–Yano (HKY) model, with different base frequencies. We used the BIONJ technique to build the starting tree and optimized the topology and branch lengths [96]. Finally, the terminal branch lengths were extracted using the Newick Utilities v.1.6 and were used as a proxy for the age of the insertions [97]. We ended up with the age estimates for 5,389 TE sequences from 116 TE families belonging to all TE orders.

We analyzed the length of the TEs by calculating the “TE length ratio (%)” defined as the length of each TE divided by the family canonical length and expressed in percentage. Then, we applied the Wilcoxon rank sum test for determining whether the distribution of the TE Length Ratio values was different between different TE classes.

### Signatures of selective sweeps

In order to detect signatures of positive selection we applied three different methods for identifying selective sweeps: *iHS* [44], *H12* [45], and *nS_L_* [46]. We separately analyzed two datasets of individually sequenced populations from Europe and North America. For the EU populations we used sequences from 158 strains belonging to four different populations: 16 strains from Castellana Grotte (Bari, South Italy) [63], 27 strains from Stockholm (Sweden) [63], 96 strains from Lyon (France) [89, 98] and 19 strains from Houten (The Netherlands) [99]. We pooled the sequences from the four European populations as it has been described that there is no evidence of latitudinal population structure in European populations [47]. This allowed us to analyse a similar number of strains in the two continents. For the Sweden and Italian populations, we first obtained the *vcf* and *bam* files from [47], we filtered out all non-SNP variants and then we used *Shapeit v2.r837* [100] for estimating haplotypes (phasing). For the French and Dutch populations we first downloaded consensus sequences from the Drosophila Genome Nexus (DGN) 1.1 [98], and we then created a *SNP-vfc* file using a custom python script. We then merged all EU populations in a single *SNP-vcf* file using *vcftools v.0.1.15* [101]. For the NA population we used the *SNP-vcf* file as provided by the Genetic Reference Panel (DGRP) for 141 strains collected in Raleigh, North Carolina [90, 91]. *iHS* was calculated using the *iHSComputer* software (https://github.com/sunthedeep/iHSComputer). We created *iHSComputer* input files (*SNPs-TEs* files) by adding the *T-lex2* information to the *SNP-vcf* file. For each TE and each strain we codified the presence/absence of the TE in a biallelic way and place them in the midpoint coordinate of the TE. Note that only presence/absence results from *T-lex2* were taken into account, leaving “polymorphic” and “no data” as missing data positions [31].

The presence of the TE was considered as the ‘derived’ state and the absence as the ‘ancestral’ state. Since *iHSComputer* runs for each chromosome separately, we created 100kbp-windows recombination files for each chromosome based on the recombination map from [87]. We standardized iHS values according to Voight *et al*. [44] and determined its significance by comparing iHS value for the TEs against the empirical distribution of iHS values for SNPs falling within the first 8-30 base pairs of small introns (<=65 bp) which are considered to be neutrally evolving [102]. Two empirical distributions were generated: one for the SNPs present at high frequency in the out-of-Africa and in the African populations, and another one for SNPs present at high frequency in out-of-Africa populations but present at low frequency in the African population (S11 Fig). TEs with *iHS* values falling outside the 5th percentile of the corresponding empirical distribution of neutral SNPs were considered significant.

The *H12* statistic was calculated using the SelectionHapStats software (https://github.com/ngarud/SelectionHapStats/, [45]. We formatted the *SNPs-TEs* files previously used in the *iHS* calculation and run the *H12_H2H1.pyscript* for each TE in the *singleWindow* mode using 100 SNPs as the window size. We first selected windows in the top 15% most extreme H12 values. We then checked whether haplotypes in these windows contained the TE in at least 50% of the strains for at least one of the three most frequent haplotypes. Only TEs that fulfil this condition were considered significant. Note that 17 out of the 18 significant TEs are present in the first or second most frequent haplotype.

The *nS*_L_ statistic was calculated using *selscan v1.1* [103]. Input files were generated based on the *SNPs-TEs* files from the *iHS* calculation. We created one *tped* file for each TE and removed all strains and positions containing missing data. Extreme *nS*L values were determined using the *norm* program for the analysis of *selscan* output. Unstandardized *nS*_L_ values were normalized in 10 frequency bins across the entire chromosome and significant *nS*_L_ values were determined using the --*crit-percent* 0.05 parameter.

### Population differentiation using FST for latitudinal distant populations

We calculate the Fixation index (Fst) between pairs of latitudinal distant populations for each of the three continents. We created *vcf* files for the TEs based on *T-lex2* results and used *vcftools v.0.1.15* [101] for calculating the pairwise FST estimator [48]. The pairwise calculations performed for each continent were: Europe: Italy vs. Sweden [63] and Vesanto vs. Nicosia [47]; Oceania: Innisfail vs. Yering [104] and Queensland vs. Tasmania [73] and North America: Maine vs. Florida [73], and Maine vs. Florida [74]. For each pair, we calculated FST values for all TEs and tested them against the empirical distribution of FST values of neutral SNPs while controlling for TE frequency in the African population [81].

Innisfail, Yering, Maine and Florida SNP callings were obtained from the Dryad Digital Repository (http://datadryad.org/resource/doi:10.5061/dryad.7440s, [74, 104]. Queensland, Tasmania, Florida and Maine SNP callings from Reinhardt *et al*. [73] were provided by Dr. Andrew Kern. Italy and Sweden SNP callings were obtained from Mateo *et al*. [63]. Vesanto and Nicosia SNP callings were obtained from Kapun et al. [47]. FST values for neutral SNPs were also calculated using *vcftools v.0.1.15* [101]. Then, for each pairwise comparison we created two empirical distributions of FST values of neutral SNPs: one for SNPs that were at low frequency in Zambia and other for SNPs that were at high frequency in Zambia. FST values of TEs at high frequency in Zambia were compared with the distribution of neutral SNPs FST at high frequency in Zambia and FST values of TEs at low frequency in Zambia were compared with the low frequency SNPs distribution. We considered a TE to be significantly differentiated when its FST value was greater than the percentile 95th of the corresponding empirical distribution.

Overall, we calculated FST values for 254 TEs in at least one pair of populations and we found 78 of them showing extreme values when comparing with the distribution of FST from neutral SNPs (S7 Table). 67 of these 78 TEs were consistently present at high frequencies in populations located in high latitudes or in low latitudes. 43 of the 67 TEs were present at high frequencies in low latitude populations in at least one pairwise comparison, and 24 TEs were present at high frequencies in high latitude populations in at least one pairwise comparison (S7 and S16 Tables). Finally, to be conservative, we only considered those TEs with significant FST values in at least two populations and always present at high frequencies in populations located in high or low latitude (concordant FST).

### TE location

We analyzed whether TEs were located at specific regions in the genome regarding the nearest gene. We used TEs and gene coordinates from FlyBase r6.04 [84, 85] and considered both coding and non-coding genes. For each TE, we determined whether it was located inside a gene or in an intergenic region. We further classify the TEs located in intergenic regions in those located at more or less than 1kb of the nearest gene. For TEs present inside a gene we further determined the class site overlapping with the TE annotation: *Exon, UTR, Intron*. If the TE is inserted in an intron, we checked whether it was inserted in the first intron, where is more likely to affect expression [105, 106].

### Expression quantitative trait loci (eQTL) analysis

We use Matrix eQTL v2.1.1 [51] to calculate correlations between the presence/absence of the TEs and the expression of nearby genes. We used expression data from the DRGP lines (Raleigh, North Carolina, [52]) as available in the DGRP2 repository (http://dgrp2.gnets.ncsu.edu/data.html) and the presence/absence TE information for the DGRP lines for which *T-lex2* was successfully run (see above). *T-lex2* identified TEs for 1,603 in the DRGP lines and 1,177 of them contain at least one gene at less than 1kb of any of the two junction coordinates of the TEs. One line (RAL-591) was not present in the expression data, so we ended up with 140 lines in the dataset. For each line, we used the average of the normalized gene expression value from the two replicates and analyzed female and male data separately. For the genotyping data, we used both the start and the end coordinates of the 1,615 TE as positions in the genome and codified the absence (0), polymorphic (1), presence (2) and no data (NA) from *T-lex2* output using a custom python script. *Matrix eQTL* was run with default parameters, applying only the *Linear* model and with a *cisDist=1000*, meaning that we considered only genes that were at less than 1kb from any of the junction coordinates of the TE. We then evaluated the significance of the correlations as provided by the *Matrix eQTL* software and we considered TEs that were significant in at least one sex. From the 1,177 analyzed TEs, we kept only the 638 TEs located at high recombination rate regions and classified them according to their frequency in the DGRP population as: HighFreq (10% < frequency < 95%), LowFreq (frequency ≤ 10%) and Fixed (frequency ≥ 95%). LowFreq TEs were further classified as Private if only one strain was containing the TE. 235 of the 300 candidate adaptive TEs were included in the 638 dataset.

### Functional enrichment analysis

We performed functional enrichment analysis for Gene Ontology (GO) biological process for the genes nearby TEs using the DAVID functional annotation cluster tool (*v.6.8*) [49, 50]. Based on TE and gene coordinates from FlyBase r6.04 [84, 85], we selected genes located at less than 1kb as the ones putatively likely affected by the TEs, since this is the approximate size of the promoter region in *D. melanogaster* [107]. If there were no genes at less than 1kb, we selected the closest one. All comparisons were performed using the full list of genes in *D. melanogaster* as the background. We considered DAVID clusters as significant when the enrichment score (ES) was higher than 1.3 as described in Huang da *et al*. [49].

In addition, in December 2016 we searched the literature using PubMed to find publications that identified genes associated with phenotypic traits studied in the DGRP project (olfactory behavior, alcohol exposure, desiccation, aggressiveness, cold tolerance, pigmentation, starvation, mating behavior, and oxidative stress). We also included phenotypic traits for which there is gene expression data available (heavy-metal stress, xenobiotic stress, diapause, locomotor behavior, and hypoxia). Finally, we looked for publications related with immunity, heat-shock stress, and circadian behavior as these three are relevant adaptive traits in Drosophila. We included genome-wide studies (GWAS, QTL, gene expression, and protein-protein interactions) and candidate-gene studies (S9 Table). We generated lists of candidate genes for each one of the 17 different fitness-related traits. We then converted the gene names to Flybase gene identifiers. This step was necessary because in *D. melanogaster* genes often have more than one name but all genes have a single Flybase identifier. To construct our final candidate gene lists, we only considered those genes that were present in two or more independent publications. We then checked whether the genes nearby the 300 HighFreq TEs, the 65 TEs with evidence of positive selection, the 174 OOA, and the 111 AF-OOA TEs were present in our candidate gene lists. We used Chi-square test to determine whether different sets of TEs showed more genes previously associated with different stress-related and behavior-related traits than expected by chance.

## Acknowledgments

We thank members of the González lab for comments on the manuscript. This work was funded by a grant from the European Commission (H2020-ERC-2014-CoG-647900) and by the Ministerio de Economia y Competitividad (BFU2014-57779-P).

## Supporting information

**S1 Fig. Distribution of number of TEs that are present at >0.10 and < 0.95 frequency by number of populations in which they are present at that frequency**. We considered TEs to be present at high frequency (HighFreq) when they fulfil the frequency condition in at least three samples (represented by blue bars in the figure).

**S2 Fig. Comparison of age estimations obtained by Bergman and Bensasson (2007) and the estimations obtained in this work. Only the 417 TEs that are common between the two studies are plotted**. A) TE age distribution of the 417 TEs based on Bergman and Bensasson (2007) and in this work. Note that there are 10 insertions that showed extreme age values in our dataset (> 0.12). B) Correlation between the two age estimates before and after removing the 10 TEs with extreme age values in our data set (n = 407).

**S3 Fig. Venn diagrams for the 36 HighFreq TEs with significant evidence of selective sweeps. A) Overlapping between TEs showing significant results for the different selective sweeps statistics (*iHS, H12* and *nSL*). B**) Overlapping between TEs showing at least one significant test in the North American (NA) and/or the European (EU) population. The percentage between brackets is regarding the total number of significant TEs (36). Numbers between square brackets show the number of TEs for which we were able to calculate at least one of the sweep statistics.

**S4 Fig. Venn diagrams showing the overlap between TEs showing significant FST values in at least one pair of populations**. A) TEs present at high frequency in populations located at low latitude locations. B) TEs present at high frequency in populations located at high latitude locations.

**S5 Fig. Functional enrichment analysis of genes nearby OOA and AF-OOA TEs. A) Significant Gene Ontology Clusters according to DAVID functional annotation tool**. Only the top six significant clusters are showed (enrichment score > 1.3). The horizontal axis represents DAVID enrichment score (see S9C and S9D Tables for details). B) Significantly overrepresented fitness-related genes according to previous genome association studies. All FDR corrected p-values < 0.05, Chi-square test (see S11C and S11D Tables for details). The horizontal axis represent the log10(χ^2^). In both cases, A) and B), numbers nearby each bar indicate total number of genes in that category. Bar colors indicates similar biological functions of the clusters (A) and the fitness-related traits (B): green: stress response; red: behavior; blue: development; and yellow: transport.

**S6 Fig. Distribution of the number of TEs (y axis) by the number of strains for which *T-lex2* estimated frequencies in the 8 individually-sequenced populations**.

**S7 Fig. Distribution of mapped reads for the presence module (red), absence module (green) and total number of reads (blue) for each one of the 48 DrosEU samples (Kapun *et al*. 2018)**.

**S8 Fig. Correlation between frequencies estimated with data obtained using different sequencing strategies in the Stockholm (Sweden) population**. Frequencies calculated using individual strain sequencing (x) (Mateo et al 2018) and pool sequencing (y). Pearson correlation coefficient r = 0.98, p-value < 2.2e^-16^.

**S9 Fig. Histogram showing the number of TEs (y axis) and the number of samples for which we were able to estimate its frequency**.

**S10 Fig. TE frequencies estimated using all strains (x axis) vs. frequencies estimated after removing strains that contain inversions (y axis) for different individually-sequenced populations. A**) Zambia (Lack et al., 2015), **B**) France (Pool et al., 2012), **C**) DGRP (Raleigh) (Huang *et al*. 2014; Mackay *et al*. 2012), **D**) Italy (Bari) and **E**) Sweden (Stockholm) (Mateo et al 2018). All Pearson correlation coefficients r=0.99 and p-value < 2.2e^-16^.

**S11 Fig. Distribution of iHS values obtained for TEs (red) and neutral SNPs (cyan) in the North American population (DGRP, Raleigh, North Carolina)**. **A**) Distribution of iHS values for all TEs and neutral SNPs. **B**) Distribution of iHS values for TEs and neutral SNPs at high frequency (> 0.10) in the OOA population (Raleigh) and in the African population (Zambia). **C**) Distribution of iHS values for TEs and neutral SNPs at high frequency (> 0.10) in the OOA population, but at low frequency in the African population.

**S1 Table. Information for the 91 samples used in this study**.

**S2 Table. Frequency estimations using Tlex2 for the 1,615 TEs at each of the 91 samples**. NA indicates that the frequency could not be estimated for that TE in the given sample. Recombination estimates according to Comeron *et al*. (2012) and Fiston-Lavier *et al*. (2010) are showed for each TE. Class column indicates the category at which each TE was classified.

**S3 Table. TEs at each category classified as young (divergence < 0.01) or old (divergence ≥ 0.01)**. P-values are from Chi-square test when comparing TEs at each category the expectations when considering All 1.615 TEs.

**S4 Table. TE Length Ratio statistics**. At the top, mean and median TE Length Ratio (%) for each TE category. At the bottom, results for the Wilcoxon rank sum test and Kruskal Wallis test among different TE categories.

**S5 Table. Number of TEs showing significant values in the selection tests for each HighFreq category**. For each sweep test (iHS, H12 and *nS*_L_), “Continent” column indicates population used for the analysis: NA: North America or EU: Europe. For each HighFreq category, table shows the number of significant TEs / number of TEs for which the test was calculated. “At least one test” indicates the number of TEs at each category showing at least one test significant / TEs with at least one test calculated.

**S6 Table. List of 36 TEs showing at least one significant (highlighted in red) selective sweep test (iHS, H12 or *nS*_L_)**.

**S7 Table. List of the 254 HighFreq TEs with at least one pairwise Fst calculation performed**. Category indicates the classification of the TE according to Figure 2. For each continent, two pairwise comparisons were performed. Values for each comparison are the Fst (in red the significant ones). Concordant Fst indicates whether TEs with significant Fst were at high frequency in the same climate zone in more than one population. Concordance information indicates, for each significant pairwise calculation (separated by ‘;’) the continent (EU, NA or OC) and the climate zone at which the TE is a higher frequency (Tropical/Mild Temperature, Snow).

**S8A Table**: Results of gene ontology (GO) enrichment test for the 83 genes nearby the 65 TEs showing evidence of selection (ES).

**S8B Table**. Results of gene ontology (GO) enrichment test for the 363 genes nearby the 300 HigFreq TEs.

**S8C Table**: Results of gene ontology (GO) enrichment test for the 215 genes nearby the 174 OOA TEs.

**S8D Table**. Results of gene ontology (GO) enrichment test for the 143 genes nearby the 111 AF-OOA TEs.

**S9 Table. Gene association studies analyzing different fitness-related phenotypes**.

**S10 Table. Enrichment of genes previously described as associated with different stress-related and behaviour-reltaed traits in the different datasets analyzed**. A) Genes the 65 TEs with evidence of selection. B) Genes nearby the 300 HighFreq TEs. C) Genes nearby the 174 OOA TEs. D) Genes nearby the 111 AF-OOA TEs.

**S11 Table. 19 TEs showing significant correlation with the expression of nearby genes**. Results are divided in correlations obtained with male and female expression data (Huang et al. 2015). beta: Effect size estimate, t-stat: Test statistic (t-statistic of T-test), p-value: p-value for the linear regression. FDR: False discovery rate estimated with Benjamini–Hochberg procedure. * TEs showing evidence of selection (Table 1, main text).

**S12 Table. Genomic location of different TE categories**. Percentages and rigth-tail p-values are showed when the Chi-square test is significant. (A) Localization of TEs regarding the nearest gene across categories. (B) Localization of intragenic TEs across TE categories.

**S13 Table. TE classes across different TE categories**. P-values and percentages are showed in bold when significant enrichment according to Chi-square test p-value < 0.05 when comparing with All TEs.

**S14 Table. Enrichment test for TE families. For each family, table shows the number of TEs at each category**. HighFreq TEs correspond to the sum of AF, AF-NA, AF-OOA and OOA. p-value (Bonf.) indicates Bonferroni corrected p-values for Chi-square test when comparing HighFreq, AF-OOA and OOA TEs against All TEs. In red p-values < 0.05.

**S15 Table. Genomic coordinates of cosmopolitan inversion (Kapun *et al*. 2016) analyzed in order to determine its influence on the transposable elements frequency calculation**.

**S16 Table. Summary statistics for the pairwise Fst calculations**. <u>TEs with Fst</u>: Number of TEs for which it was possible to calculate Fst. <u>Signif. (Africa H/L)</u>: Total number of significant TEs. Between brackets: H: Number of significant TEs identified using the distribution of neutral SNPs that are at high frequency in Africa. L: Number of significant TEs identified using the distribution of neutral SNPs that are at low frequency in Africa (see Material and Methods). <u>Low Latitude (HighFreq)</u>: Significant TEs that are at high recombination rate regions (HRR) and are at high frequency only in populations located in low latutidinal regions. <u>High Latitude (HighFreq)</u>: Significant TEs that are at high recombination rate regions (HRR) and are at high frequency in populations located in high latutidinal regions. <u>Both (HighFreq)</u>: Significant TEs that are at high recombination rate regions (HRR) and are at high frequency in populations from both, low and high latutidinal regions.

## References

1. Pardo-Diaz C, Salazar C, Jiggins CD. Towards the identification of the loci of adaptive evolution. Methods Ecol Evol. 2015;6(4):445–64. doi: 10.1111/2041-210X.12324. PubMed PMID: 25937885; PubMed Central PMCID: PMCPMC4409029.

2. Hoban S, Kelley JL, Lotterhos KE, Antolin MF, Bradburd G, Lowry DB, et al. Finding the Genomic Basis of Local Adaptation: Pitfalls, Practical Solutions, and Future Directions. Am Nat. 2016;188(4):379–97. Epub 2016/09/14. doi: 10.1086/688018. PubMed PMID: 27622873; PubMed Central PMCID: PMCPMC5457800.

3. Francois O, Martins H, Caye K, Schoville SD. Controlling false discoveries in genome scans for selection. Mol Ecol. 2016;25(2):454–69. Epub 2015/12/17. doi: 10.1111/mec.13513. PubMed PMID: 26671840.

4. Fan S, Hansen ME, Lo Y, Tishkoff SA. Going global by adapting local: A review of recent human adaptation. Science. 2016;354(6308):54–9. doi: 10.1126/science.aaf5098. PubMed PMID: 27846491; PubMed Central PMCID: PMCPMC5154245.

5. Jeong C, Di Rienzo A. Adaptations to local environments in modern human populations. Current opinion in genetics & development. 2014;29:1–8. Epub 2014/08/19. doi: 10.1016/j.gde.2014.06.011. PubMed PMID: 25129844; PubMed Central PMCID: PMCPMC4258478.

6. Flood PJ, Hancock AM. The genomic basis of adaptation in plants. Curr Opin Plant Biol. 2017;36:88–94. doi: 10.1016/j.pbi.2017.02.003. PubMed PMID: 28242535.

7. Gonzalez J, Lenkov K, Lipatov M, Macpherson JM, Petrov DA. High rate of recent transposable element-induced adaptation in Drosophila melanogaster. PLoS Biol. 2008;6(10):e251. doi: 10.1371/journal.pbio.0060251. PubMed PMID: 18942889; PubMed Central PMCID: PMCPMC2570423.

8. Gonzalez J, Karasov TL, Messer PW, Petrov DA. Genome-wide patterns of adaptation to temperate environments associated with transposable elements in Drosophila. PLoS Genet. 2010;6(4):e1000905. Epub 2010/04/14. doi: 10.1371/journal.pgen.1000905. PubMed PMID: 20386746; PubMed Central PMCID: PMCPMC2851572.

9. Dennis MY, Eichler EE. Human adaptation and evolution by segmental duplication. Current opinion in genetics & development. 2016;41:44–52. Epub 2016/09/02. doi:10.1016/j.gde.2016.08.001. PubMed PMID: 27584858; PubMed Central PMCID: PMCPMC5161654.

10. Rishishwar L, Wang L, Clayton EA, Marino-Ramirez L, McDonald JF, Jordan IK. Population and clinical genetics of human transposable elements in the (post) genomic era. Mobile genetic elements. 2017;7(1):1–20. Epub 2017/02/24. doi: 10.1080/2159256x.2017.1280116. PubMed PMID: 28228978; PubMed Central PMCID: PMCPMC5305044.

11. Chuong EB, Elde NC, Feschotte C. Regulatory activities of transposable elements: from conflicts to benefits. Nat Rev Genet. 2017;18(2):71–86. Epub 2016/11/22. doi: 10.1038/nrg.2016.139. PubMed PMID: 27867194; PubMed Central PMCID: PMCPMC5498291.

12. Casacuberta E, Gonzalez J. The impact of transposable elements in environmental adaptation. Mol Ecol. 2013;22(6):1503–17. Epub 2013/01/09. doi: 10.1111/mec.12170. PubMed PMID: 23293987.

13. Hua-Van A, Le Rouzic A, Boutin TS, Filee J, Capy P. The struggle for life of the genome’s selfish architects. Biol Direct. 2011;6:19. Epub 2011/03/19. doi: 10.1186/1745-6150-6-19. PubMed PMID: 21414203; PubMed Central PMCID: PMCPMC3072357.

14. Guio L, Gonzalez J. New insights on the evolution of genome content: population dynamics of transposable elements in flies and humans. Methods in Molecular Biologyin press.

15. Elbarbary RA, Lucas BA, Maquat LE. Retrotransposons as regulators of gene expression. Science. 2016;351(6274):aac7247. Epub 2016/02/26. doi: 10.1126/science.aac7247. PubMed PMID: 26912865; PubMed Central PMCID: PMCPMC4788378.

16. Petrov DA, Fiston-Lavier AS, Lipatov M, Lenkov K, Gonzalez J. Population genomics of transposable elements in Drosophila melanogaster. Mol Biol Evol. 2011;28(5):1633–44. doi: 10.1093/molbev/msq337. PubMed PMID: 21172826; PubMed Central PMCID: PMCPMC3080135.

17. Chuong EB, Elde NC, Feschotte C. Regulatory evolution of innate immunity through co-option of endogenous retroviruses. Science. 2016;351(6277):1083–7. Epub 2016/03/05. doi: 10.1126/science.aad5497. PubMed PMID: 26941318; PubMed Central PMCID: PMCPMC4887275.

18. Daborn PJ, Yen JL, Bogwitz MR, Le Goff G, Feil E, Jeffers S, et al. A single p450 allele associated with insecticide resistance in Drosophila. Science. 2002;297(5590):2253–6. doi: 10.1126/science.1074170. PubMed PMID: 12351787.

19. Chung H, Bogwitz MR, McCart C, Andrianopoulos A, Ffrench-Constant RH, Batterham P, et al. Cis-regulatory elements in the Accord retrotransposon result in tissue-specific expression of the Drosophila melanogaster insecticide resistance gene Cyp6g1. Genetics. 2007;175(3):1071–7. Epub 2006/12/21. doi: 10.1534/genetics.106.066597. PubMed PMID: 17179088; PubMed Central PMCID: PMCPMC1840086.

20. Kuhn A, Ong YM, Cheng CY, Wong TY, Quake SR, Burkholder WF. Linkage disequilibrium and signatures of positive selection around LINE-1 retrotransposons in the human genome. Proc Natl Acad Sci U S A. 2014;111(22):8131–6. Epub 2014/05/23. doi: 10.1073/pnas.1401532111. PubMed PMID: 24847061; PubMed Central PMCID: PMCPMC4050588.

21. Blumenstiel JP, Chen X, He M, Bergman CM. An age-of-allele test of neutrality for transposable element insertions. Genetics. 2014;196(2):523–38. Epub 2013/12/18. doi: 10.1534/genetics.113.158147. PubMed PMID: 24336751; PubMed Central PMCID: PMCPMC3914624.

22. Quadrana L, Bortolini Silveira A, Mayhew GF, LeBlanc C, Martienssen RA, Jeddeloh JA, et al. The Arabidopsis thaliana mobilome and its impact at the species level. Elife. 2016;5. Epub 2016/06/04. doi: 10.7554/eLife.15716. PubMed PMID: 27258693; PubMed Central PMCID: PMCPMC4917339.

23. Stuart T, Eichten SR, Cahn J, Karpievitch YV, Borevitz JO, Lister R. Population scale mapping of transposable element diversity reveals links to gene regulation and epigenomic variation. Elife. 2016;5. Epub 2016/12/03. doi: 10.7554/eLife.20777. PubMed PMID: 27911260; PubMed Central PMCID: PMCPMC5167521.

24. Gonzalez J, Petrov DA. Evolution of genome content: population dynamics of transposable elements in flies and humans. Methods in molecular biology (Clifton, NJ). 2012;855:361–83. Epub 2012/03/13. doi: 10.1007/978-1-61779-582-4_13. PubMed PMID: 22407716.

25. David JR, Capy P. Genetic variation of Drosophila melanogaster natural populations. Trends Genet. 1988;4(4):106–11. Epub 1988/04/01. PubMed PMID: 3149056.

26. Li H, Stephan W. Inferring the demographic history and rate of adaptive substitution in Drosophila. PLoS Genet. 2006;2(10):e166. doi: 10.1371/journal.pgen.0020166. PubMed PMID: 17040129; PubMed Central PMCID: PMCPMC1599771.

27. Hervas S, Sanz E, Casillas S, Pool JE, Barbadilla A. PopFly: the Drosophila population genomics browser. Bioinformatics. 2017;33(17):2779–80. doi: 10.1093/bioinformatics/btx301.

28. Gramates LS, Marygold SJ, Santos Gd, Urbano J-M, Antonazzo G, Matthews BB, et al. FlyBase at 25: looking to the future. Nucleic Acids Research. 2017;45(D1):D663–D71. doi: 10.1093/nar/gkw1016.

29. Kaminker JS, Bergman CM, Kronmiller B, Carlson J, Svirskas R, Patel S, et al. The transposable elements of the Drosophila melanogaster euchromatin: a genomics perspective. Genome Biol. 2002;3(12):Research0084. Epub 2003/01/23. PubMed PMID: 12537573; PubMed Central PMCID: PMCPMC151186.

30. Quesneville H, Bergman CM, Andrieu O, Autard D, Nouaud D, Ashburner M, et al. Combined Evidence Annotation of Transposable Elements in Genome Sequences. PLOS Computational Biology. 2005;1(2):e22. doi: 10.1371/journal.pcbi.0010022.

31. Fiston-Lavier AS, Barron MG, Petrov DA, Gonzalez J. T-lex2: genotyping, frequency estimation and re-annotation of transposable elements using single or pooled next-generation sequencing data. Nucleic Acids Res. 2015;43(4):e22. doi: 10.1093/nar/gku1250. PubMed PMID: 25510498; PubMed Central PMCID: PMCPMC4344482.

32. Chen D, Chen HW. Using the Köppen classification to quantify climate variation and change: An example for 1901–2010. Environmental Development. 2013;6:69–79. doi: http://dx.doi.org/10.1016/j.envdev.2013.03.007.

33. Hill WG, Robertson A. The effect of linkage on limits to artificial selection. Genetical research. 1966;8(3):269–94. Epub 1966/12/01. PubMed PMID: 5980116.

34. Smith JM, Haigh J. The hitch-hiking effect of a favourable gene. Genetical research. 1974;23(1):23–35. Epub 1974/02/01. PubMed PMID: 4407212.

35. Charlesworth B, Morgan MT, Charlesworth D. The effect of deleterious mutations on neutral molecular variation. Genetics. 1993;134(4):1289–303. Epub 1993/08/01. PubMed PMID: 8375663; PubMed Central PMCID: PMCPMC1205596.

36. Hudson RR, Kaplan NL. Deleterious background selection with recombination. Genetics. 1995;141(4):1605–17. Epub 1995/12/01. PubMed PMID: 8601498; PubMed Central PMCID: PMCPMC1206891.

37. Barron MG, Fiston-Lavier AS, Petrov DA, Gonzalez J. Population genomics of transposable elements in Drosophila. Annu Rev Genet. 2014;48:561–81. doi: 10.1146/annurev-genet-120213-092359. PubMed PMID: 25292358.

38. Castellano D, Coronado-Zamora M, Campos JL, Barbadilla A, Eyre-Walker A. Adaptive Evolution Is Substantially Impeded by Hill–Robertson Interference in Drosophila. Molecular Biology and Evolution. 2016;33(2):442–55. doi: 10.1093/molbev/msv236. PubMed PMID: PMC4794616.

39. Sellis D, Callahan BJ, Petrov DA, Messer PW. Heterozygote advantage as a natural consequence of adaptation in diploids. Proceedings of the National Academy of Sciences of the United States of America. 2011;108(51):20666–71. doi: 10.1073/pnas.1114573108. PubMed PMID: PMC3251125.

40. Bergman CM, Bensasson D. Recent LTR retrotransposon insertion contrasts with waves of non-LTR insertion since speciation in <em>Drosophila melanogaster</em>. Proceedings of the National Academy of Sciences. 2007;104(27):11340–5. doi: 10.1073/pnas.0702552104.

41. Montgomery E, Charlesworth B, Langley CH. A test for the role of natural selection in the stabilization of transposable element copy number in a population of Drosophila melanogaster. Genetical research. 1987;49(1):31–41. Epub 1987/02/01. PubMed PMID: 3032743.

42. Petrov DA, Aminetzach YT, Davis JC, Bensasson D, Hirsh AE. Size matters: non-LTR retrotransposable elements and ectopic recombination in Drosophila. Mol Biol Evol. 2003;20(6):880–92. Epub 2003/04/30. doi: 10.1093/molbev/msg102. PubMed PMID: 12716993.

43. Petrov DA, Chao YC, Stephenson EC, Hartl DL. Pseudogene evolution in Drosophila suggests a high rate of DNA loss. Mol Biol Evol. 1998;15(11):1562–7. Epub 2003/02/08. PubMed PMID: 12572619.

44. Voight BF, Kudaravalli S, Wen X, Pritchard JK. A Map of Recent Positive Selection in the Human Genome. PLOS Biology. 2006;4(3):e72. doi: 10.1371/journal.pbio.0040072.

45. Garud NR, Messer PW, Buzbas EO, Petrov DA. Recent Selective Sweeps in North American Drosophila melanogaster Show Signatures of Soft Sweeps. PLOS Genetics. 2015;11(2):e1005004. doi: 10.1371/journal.pgen.1005004.

46. Ferrer-Admetlla A, Liang M, Korneliussen T, Nielsen R. On detecting incomplete soft or hard selective sweeps using haplotype structure. Mol Biol Evol. 2014;31(5):1275–91. Epub 2014/02/21. doi: 10.1093/molbev/msu077. PubMed PMID: 24554778; PubMed Central PMCID: PMCPMC3995338.

47. Kapun M, Barron Aduriz MG, Staubach F, Vieira J, Obbard D, Goubert C, et al. Genomic analysis of European Drosophila populations reveals longitudinal structure and continent-wide selection. bioRxiv. 2018. doi: 10.1101/313759.

48. Weir BS, Cockerham CC. Estimating F-statistics for the analysis of population structure. Evolution. 1984;38(6):1358–70. doi: 10.1111/j.1558-5646.1984.tb05657.x. PubMed PMID: 28563791.

49. Huang da W, Sherman BT, Lempicki RA. Bioinformatics enrichment tools: paths toward the comprehensive functional analysis of large gene lists. Nucleic Acids Res. 2009;37(1):1–13. Epub 2008/11/27. doi: 10.1093/nar/gkn923. PubMed PMID: 19033363; PubMed Central PMCID: PMCPMC2615629.

50. Huang DW, Sherman BT, Lempicki RA. Systematic and integrative analysis of large gene lists using DAVID bioinformatics resources. Nat Protocols. 2008;4(1):44–57. doi: http://www.nature.com/nprot/iournal/v4/n1/suppinfo/nprot.2008.211S1.html.

51. Shabalin AA. Matrix eQTL: ultra fast eQTL analysis via large matrix operations. Bioinformatics. 2012;28(10):1353–8. doi: 10.1093/bioinformatics/bts163. PubMed PMID: PMC3348564.

52. Huang W, Carbone MA, Magwire MM, Peiffer JA, Lyman RF, Stone EA, et al. Genetic basis of transcriptome diversity in Drosophila melanogaster. Proc Natl Acad Sci U S A. 2015;112(44):e6010–9. Epub 2015/10/21. doi: 10.1073/pnas.1519159112. PubMed PMID: 26483487; PubMed Central PMCID: PMCPMC4640795.

53. Mackay TF. Mutations and quantitative genetic variation: lessons from Drosophila. Philosophical transactions of the Royal Society of London Series B, Biological sciences. 2010;365(1544):1229–39. Epub 2010/03/24. doi: 10.1098/rstb.2009.0315. PubMed PMID: 20308098; PubMed Central PMCID: PMCPMC2871822.

54. Cridland JM, Thornton KR, Long AD. Gene Expression Variation in Drosophila melanogaster Due to Rare Transposable Element Insertion Alleles of Large Effect. Genetics. 2015;199(1):85–93. doi: 10.1534/genetics.114.170837. PubMed PMID: PMC4286695.

55. Aminetzach YT, Macpherson JM, Petrov DA. Pesticide resistance via transposition-mediated adaptive gene truncation in Drosophila. Science. 2005;309(5735):764–7. Epub 2005/07/30. doi: 10.1126/science.1112699. PubMed PMID: 16051794.

56. Guio L, Barron MG, Gonzalez J. The transposable element Bari-Jheh mediates oxidative stress response in Drosophila. Mol Ecol. 2014;23(8):2020–30. doi: 10.1111/mec.12711. PubMed PMID: 24629106.

57. Mateo L, Ullastres A, Gonzalez J. A transposable element insertion confers xenobiotic resistance in Drosophila. PLoS Genet. 2014;10(8):e1004560. doi: 10.1371/journal.pgen.1004560. PubMed PMID: 25122208; PubMed Central PMCID: PMCPMC4133159.

58. Ullastres A, Petit N, Gonzalez J. Exploring the phenotypic space and the evolutionary history of a natural mutation in Drosophila melanogaster. Mol Biol Evol. 2015;32(7):1800–14. doi: 10.1093/molbev/msv061. PubMed PMID: 25862139; PubMed Central PMCID: PMCPMC4476160.

59. Merenciano M, Ullastres A, de Cara MA, Barron MG, Gonzalez J. Multiple independent retroelement insertions in the promoter of a stress response gene have variable molecular and functional effects in Drosophila. PLoS Genet. 2016;12(8):e1006249. doi: 10.1371/journal.pgen.1006249. PubMed PMID: 27517860; PubMed Central PMCID: PMCPMC4982627.

60. Schmidt JM, Good RT, Appleton B, Sherrard J, Raymant GC, Bogwitz MR, et al. Copy number variation and transposable elements feature in recent, ongoing adaptation at the Cyp6g1 locus. PLoS Genet. 2010;6(6):e1000998. Epub 2010/06/30. doi: 10.1371/journal.pgen.1000998. PubMed PMID: 20585622; PubMed Central PMCID: PMCPMC2891717.

61. Kofler R, Betancourt AJ, Schlotterer C. Sequencing of pooled DNA samples (Pool-Seq) uncovers complex dynamics of transposable element insertions in Drosophila melanogaster. PLoS Genet. 2012;8(1):e1002487. Epub 2012/02/01. doi: 10.1371/journal.pgen.1002487. PubMed PMID: 22291611; PubMed Central PMCID: PMCPMC3266889.

62. Zhu CT, Chang C, Reenan RA, Helfand SL. Indy gene variation in natural populations confers fitness advantage and life span extension through transposon insertion. Aging. 2014;6(1):58–69. Epub 2014/02/13. doi: 10.18632/aging.100634. PubMed PMID: 24519859; PubMed Central PMCID: PMCPMC3927810.

63. Mateo L, Rech G, Gonzalez J. Genome-wide patterns of local adaptation in Drosophila melanogaster: adding intra European variability to the map. bioRxiv. 2018. doi: 10.1101/269332.

64. Villanueva-Cañas JL, Rech GE, de Cara MAR, González J. Beyond SNPs: how to detect selection on transposable element insertions. Methods in Ecology and Evolution. 2017;8(6):728–37. doi: 10.1111/2041-210X.12781.

65. Pritchard JK, Di Rienzo A. Adaptation – not by sweeps alone. Nature Reviews Genetics. 2010;11:665. doi: 10.1038/nrg2880.

66. Coop G, Witonsky D, Di Rienzo A, Pritchard JK. Using Environmental Correlations to Identify Loci Underlying Local Adaptation. Genetics. 2010;185(4):1411–23. doi: 10.1534/genetics.110.114819. PubMed PMID: PMC2927766.

67. Frichot E, François O, O’Meara B. LEA: An R package for landscape and ecological association studies. Methods in Ecology and Evolution. 2015;6(8):925–9. doi: doi:10.1111/2041-210X.12382.

68. Gautier M. Genome-Wide Scan for Adaptive Divergence and Association with Population-Specific Covariates. Genetics. 2015;201(4):1555–79. Epub 2015/10/21. doi: 10.1534/genetics.115.181453. PubMed PMID: 26482796; PubMed Central PMCID: PMCPMC4676524.

69. Ahrens CW, Rymer PD, Stow A, Bragg J, Dillon S, Umbers KDL, et al. The search for loci under selection: trends, biases and progress. Mol Ecol. 2018;27(6):1342–56. Epub 2018/03/11. doi: 10.1111/mec.14549. PubMed PMID: 29524276.

70. Zonato V, Collins L, Pegoraro M, Tauber E, Kyriacou CP. Is diapause an ancient adaptation in Drosophila? J Insect Physiol. 2017;98:267–74. doi: 10.1016/j.jinsphys.2017.01.017. PubMed PMID: 28161445.

71. Kolaczkowski B, Kern AD, Holloway AK, Begun DJ. Genomic Differentiation Between Temperate and Tropical Australian Populations of <em>Drosophila melanogaster</em>. Genetics. 2011;187(1):245–60. doi: 10.1534/genetics.110.123059.

72. Fabian DK, Kapun M, Nolte V, Kofler R, Schmidt PS, Schlotterer C, et al. Genome-wide patterns of latitudinal differentiation among populations of Drosophila melanogaster from North America. Mol Ecol. 2012;21(19):4748–69. doi: 10.1111/j.1365-294X.2012.05731.x. PubMed PMID: 22913798; PubMed Central PMCID: PMCPMC3482935.

73. Reinhardt JA, Kolaczkowski B, Jones CD, Begun DJ, Kern AD. Parallel geographic variation in Drosophila melanogaster. Genetics. 2014;197(1):361–73. doi: 10.1534/genetics.114.161463. PubMed PMID: 24610860; PubMed Central PMCID: PMCPMC4012493.

74. Bergland AO, Behrman EL, O’Brien KR, Schmidt PS, Petrov DA. Genomic evidence of rapid and stable adaptive oscillations over seasonal time scales in Drosophila. PLoS Genet. 2014;10(11):e1004775. doi: 10.1371/journal.pgen.1004775. PubMed PMID: 25375361; PubMed Central PMCID: PMCPMC4222749.

75. Kapun M, Fabian DK, Goudet J, Flatt T. Genomic evidence for adaptive inversion clines in Drosophila melanogaster. Mol Biol Evol. 2016;33(5):1317–36. Epub 2016/01/23. doi: 10.1093/molbev/msw016. PubMed PMID: 26796550.

76. Pool JE, Braun DT, Lack JB. Parallel Evolution of Cold Tolerance within Drosophila melanogaster. Mol Biol Evol. 2017;34(2):349–60. Epub 2016/10/26. doi: 10.1093/molbev/msw232. PubMed PMID: 27777283; PubMed Central PMCID: PMCPMC5526443.

77. Rahman R, Chirn GW, Kanodia A, Sytnikova YA, Brembs B, Bergman CM, et al. Unique transposon landscapes are pervasive across Drosophila melanogaster genomes. Nucleic Acids Res. 2015;43(22):10655–72. Epub 2015/11/19. doi: 10.1093/nar/gkv1193. PubMed PMID: 26578579; PubMed Central PMCID: PMCPMC4678822.

78. Nelson MG, Linheiro RS, Bergman CM. McClintock: An Integrated Pipeline for Detecting Transposable Element Insertions in Whole-Genome Shotgun Sequencing Data. G3 (Bethesda). 2017;7(8):2763–78. Epub 2017/06/24. doi: 10.1534/g3.117.043893. PubMed PMID: 28637810; PubMed Central PMCID: PMCPMC5555480.

79. Chakraborty M, VanKuren NW, Zhao R, Zhang X, Kalsow S, Emerson JJ. Hidden genetic variation shapes the structure of functional elements in Drosophila. Nature Genetics. 2018;50(1):20–5. doi: 10.1038/s41588-017-0010-y.

80. Ullastres A, Gonzalez J. in prep.

81. Lack JB, Cardeno CM, Crepeau MW, Taylor W, Corbett-Detig RB, Stevens KA, et al. The Drosophila genome nexus: a population genomic resource of 623 Drosophila melanogaster genomes, including 197 from a single ancestral range population. Genetics. 2015;199(4):1229–41. Epub 2015/01/30. doi: 10.1534/genetics.115.174664. PubMed PMID: 25631317; PubMed Central PMCID: PMCPMC4391556.

82. Campo D, Lehmann K, Fjeldsted C, Souaiaia T, Kao J, Nuzhdin SV. Whole-genome sequencing of two North American Drosophila melanogaster populations reveals genetic differentiation and positive selection. Mol Ecol. 2013;22(20):5084–97. Epub 2013/10/10. doi: 10.1111/mec.12468. PubMed PMID: 24102956; PubMed Central PMCID: PMCPMC3800041.

83. Bastide H, Betancourt A, Nolte V, Tobler R, Stöbe P, Futschik A, et al. A Genome-Wide, Fine-Scale Map of Natural Pigmentation Variation in Drosophila melanogaster. PLOS Genetics. 2013;9(6):e1003534. doi: 10.1371/journal.pgen.1003534.

84. Attrill H, Falls K, Goodman JL, Millburn GH, Antonazzo G, Rey AJ, et al. FlyBase: establishing a gene group resource for Drosophila melanogaster. Nucleic Acids Res. 2016;44(D1):D786–92. doi: 10.1093/nar/gkv1046. PubMed PMID: 26467478; PubMed Central PMCID: PMCPMC4702782.

85. Mohr SE, Hu Y, Kim K, Housden BE, Perrimon N. Resources for functional genomics studies in Drosophila melanogaster. Genetics. 2014;197(1):1–18. doi: 10.1534/genetics.113.154344. PubMed PMID: 24653003; PubMed Central PMCID: PMCPMC4012471.

86. Kapitonov VV, Jurka J. Molecular paleontology of transposable elements in the Drosophila melanogaster genome. Proc Natl Acad Sci U S A. 2003;100(11):6569–74. Epub 2003/05/14. doi: 10.1073/pnas.0732024100. PubMed PMID: 12743378; PubMed Central PMCID: PMCPMC164487.

87. Comeron JM, Ratnappan R, Bailin S. The Many Landscapes of Recombination in Drosophila melanogaster. PLOS Genetics. 2012;8(10):e1002905. doi: 10.1371/journal.pgen.1002905.

88. Fiston-Lavier AS, Singh ND, Lipatov M, Petrov DA. Drosophila melanogaster recombination rate calculator. Gene. 2010;463(1–2):18–20. doi: 10.1016/j.gene.2010.04.015. PubMed PMID: 20452408.

89. Pool JE, Corbett-Detig RB, Sugino RP, Stevens KA, Cardeno CM, Crepeau MW, et al. Population genomics of sub-saharan Drosophila melanogaster: African diversity and non-African admixture. PLoS Genet. 2012;8(12):e1003080. doi: 10.1371/journal.pgen.1003080. PubMed PMID: 23284287; PubMed Central PMCID: PMCPMC3527209.

90. Huang W, Massouras A, Inoue Y, Peiffer J, Ramia M, Tarone AM, et al. Natural variation in genome architecture among 205 Drosophila melanogaster Genetic Reference Panel lines. Genome Res. 2014;24(7):1193–208. doi: 10.1101/gr.171546.113. PubMed PMID: 24714809; PubMed Central PMCID: PMCPMC4079974.

91. Mackay TF, Richards S, Stone EA, Barbadilla A, Ayroles JF, Zhu D, et al. The Drosophila melanogaster Genetic Reference Panel. Nature. 2012;482(7384):173–8. doi: 10.1038/nature10811. PubMed PMID: 22318601; PubMed Central PMCID: PMCPMC3683990.

92. Flutre T, Duprat E, Feuillet C, Quesneville H. Considering Transposable Element Diversification in De Novo Annotation Approaches. PLOS ONE. 2011;6(1):e16526. doi: 10.1371/journal.pone.0016526.

93. Hoede C, Arnoux S, Moisset M, Chaumier T, Inizan O, Jamilloux V, et al. PASTEC: an automatic transposable element classification tool. PLoS One. 2014;9(5):e91929. Epub 2014/05/03. doi: 10.1371/journal.pone.0091929. PubMed PMID: 24786468; PubMed Central PMCID: PMCPMC4008368.

94. Thompson JD, Gibson TJ, Plewniak F, Jeanmougin F, Higgins DG. The CLUSTAL_X windows interface: flexible strategies for multiple sequence alignment aided by quality analysis tools. Nucleic Acids Res. 1997;25(24):4876–82. Epub 1998/02/28. PubMed PMID: 9396791; PubMed Central PMCID: PMCPMC147148.

95. Katoh K, Misawa K, Kuma Ki, Miyata T. MAFFT: a novel method for rapid multiple sequence alignment based on fast Fourier transform. Nucleic Acids Research. 2002;30(14):3059–66. doi: 10.1093/nar/gkf436.

96. Guindon S, Gascuel O. A simple, fast, and accurate algorithm to estimate large phylogenies by maximum likelihood. Systematic biology. 2003;52(5):696–704. Epub 2003/10/08. PubMed PMID: 14530136.

97. Junier T, Zdobnov EM. The Newick utilities: high-throughput phylogenetic tree processing in the UNIX shell. Bioinformatics. 2010;26(13):1669–70. Epub 2010/05/18. doi: 10.1093/bioinformatics/btq243. PubMed PMID: 20472542; PubMed Central PMCID: PMCPMC2887050.

98. Lack JB, Lange JD, Tang AD, Corbett-Detig RB, Pool JE. A thousand fly genomes: an expanded Drosophila genome nexus. Mol Biol Evol. 2016;33(12):3308–13. doi: 10.1093/molbev/msw195. PubMed PMID: 27687565; PubMed Central PMCID: PMCPMC5100052.

99. Grenier JK, Arguello JR, Moreira MC, Gottipati S, Mohammed J, Hackett SR, et al. Global diversity lines - a five-continent reference panel of sequenced Drosophila melanogaster strains. G3 (Bethesda). 2015;5(4):593–603. Epub 2015/02/13. doi: 10.1534/g3.114.015883. PubMed PMID: 25673134; PubMed Central PMCID: PMCPMC4390575.

100. Delaneau O, Marchini J, Zagury JF. A linear complexity phasing method for thousands of genomes. Nature methods. 2011;9(2):179–81. Epub 2011/12/06. doi: 10.1038/nmeth.1785. PubMed PMID: 22138821.

101. Danecek P, Auton A, Abecasis G, Albers CA, Banks E, DePristo MA, et al. The variant call format and VCFtools. Bioinformatics. 2011;27(15):2156–8. doi: 10.1093/bioinformatics/btr330. PubMed PMID: 21653522; PubMed Central PMCID: PMCPMC3137218.

102. Parsch J, Novozhilov S, Saminadin-Peter SS, Wong KM, Andolfatto P. On the utility of short intron sequences as a reference for the detection of positive and negative selection in Drosophila. Mol Biol Evol. 2010;27(6):1226–34. doi: 10.1093/molbev/msq046. PubMed PMID: 20150340; PubMed Central PMCID: PMCPMC2877998.

103. Szpiech ZA, Hernandez RD. selscan: An Efficient Multithreaded Program to Perform EHH-Based Scans for Positive Selection. Molecular Biology and Evolution. 2014;31(10):2824–7. doi: 10.1093/molbev/msu211.

104. Bergland AO, Tobler R, Gonzalez J, Schmidt P, Petrov D. Secondary contact and local adaptation contribute to genome-wide patterns of clinal variation in Drosophila melanogaster. Mol Ecol. 2016;25(5):1157–74. doi: 10.1111/mec.13455. PubMed PMID: 26547394; PubMed Central PMCID: PMCPMC5089930.

105. Chen D, Liang Z. The Analysis of Sequence Features of Introns with Drosophila RP genes. International Journal of Information Processing and Management (IJIPM). 2013;4(1):6. doi: doi:10.4156/ijipm.vol4.issue1.6.

106. Park SG, Hannenhalli S, Choi SS. Conservation in first introns is positively associated with the number of exons within genes and the presence of regulatory epigenetic signals. BMC Genomics. 2014;15(1):526. doi: 10.1186/1471-2164-15-526. PubMed PMID: PMC4085337.

107. Hoskins RA, Carlson JW, Wan KH, Park S, Mendez I, Galle SE, et al. The Release 6 reference sequence of the Drosophila melanogaster genome. Genome Res. 2015;25(3):445–58. Epub 2015/01/16. doi: 10.1101/gr.185579.114. PubMed PMID: 25589440; PubMed Central PMCID: PMCPMC4352887.

